# HOXB13 suppresses *de novo* lipogenesis through HDAC3-mediated epigenetic reprogramming

**DOI:** 10.1101/2021.10.04.463081

**Authors:** Xiaodong Lu, Ka-wing Fong, Fang Wang, Galina Gritsina, Sylvan C. Baca, Jacob E. Berchuck, Jenny Ross, Eva Corey, Navdeep Chandel, William J. Catalona, Ximing Yang, Matthew L. Freedman, Jonathan C. Zhao, Jindan Yu

## Abstract

HOXB13, a homeodomain transcription factor, critically regulates androgen receptor (AR) function and promotes androgen-dependent prostate cancer (PCa) growth. However, the functions of HOXB13 in an AR-independent context remain elusive. Here we report an essential role of HOXB13 in directly suppressing lipogenic transcriptional programs in both AR-positive and -negative PCa cells. The MEIS domain (aa70-150) of HOXB13 interacts with the histone deacetylase HDAC3, which is disrupted by HOXB13 G84E mutation that has been associated with early-onset PCa. Thus, HOXB13 wildtype (WT), but not G84E mutant, recruits HDAC3 to lipogenic enhancers to catalyze histone de-acetylation and suppress lipogenic programs. HOXB13 knockdown unleashes the expression of key lipogenic regulators such as fatty acid synthase (FASN), requiring HDAC3. Analysis of human tissues revealed that HOXB13 is lost in about 30% of metastatic castration-resistant PCa, at least in part, through DNA hypermethylation. Functionally, loss of HOXB13 leads to massive lipid accumulation in PCa cells, thereby promoting cell motility *in vitro* and fueling xenograft tumor metastasis *in vivo*, which is mitigated by pharmaceutical inhibitors of FASN. In summary, our study discovers an essential AR-independent function of HOXB13 in repressing *de novo* lipogenesis and inhibiting tumor metastasis and defines a subclass of PCa that may benefit from lipogenic pathway inhibitors.

## INTRODUCTION

Recent evidences have suggested dysregulation of lipid metabolism as a hallmark of prostate cancer (PCa) progression^1–3^. Compared to benign tissues, PCa is marked by increased expression of critical enzymes of the lipogenic pathway, including the fatty acid synthase (FASN) that catalyzes all of the reactions of *de novo* lipogenesis generating palmitic acid and the sterol regulatory element-binding proteins (SREBPs, also called SREBFs) that bind to the promoters of most enzymes involved in sterol biosynthesis^4, 5^. Androgen receptor (AR), a key driver of PCa development, is a major regulator of lipid metabolism, controlling the expression of more than 20 enzymes involved in the synthesis, uptake, and metabolism of lipids ^6^. Recently, PTEN loss, MAPK activation, and nuclear pyruvate dehydrogenase complex have also been shown to activate SREBP-mediated transcriptional regulation of lipid biosynthesis to promote prostate tumorigenesis^2, 3, 7^. Accordingly, accumulation of lipid droplets has been observed in aggressive clinical prostate tumors and metastatic deposits, as well as in circulating prostate tumor cells^8^. Androgens stimulate lipid accumulation in PCa cells, which is abolished by AR antagonists^9, 10^. Reactivation of AR-induced lipid biosynthesis has been reported to drive metastatic castration-resistant prostate cancer (mCRPC)^11^, while inhibition of *de novo* lipogenesis reduces mCRPC growth ^4^. Thus, the mechanisms driving lipid metabolism in mCRPC are of broad importance in understanding the progression of the disease and identify critical targets for therapeutic intervention.

HOXB13 is a member of the homeobox (HOX) family transcription factors that recognize and bind conserved DNA motifs^12, 13^. HOXB13 is exclusively expressed mainly in the adult prostate and at a much lower level in the colorectum^14, 15^. Understanding of HOXB13 molecular function in PCa have largely limited to its interaction with AR protein and regulation of AR cistrome. HOXB13 has been shown to have multifaceted roles in potentially initiating, tethering, or antagonizing AR binding to chromatin depending on the genomic loci^16^. Consistent with its prostate-specific expression, transgenic mice studies have shown that HOXB13 expression is required for epithelial differentiation of the ventral prostate^17^. However, in these mice HOXB13 knockout does not have a global effect on AR signaling. When expressed in benign prostate cells along with FOXA1, HOXB13 facilitates the reprogramming of AR to a PCa-specific cistrome^18^. However, another study reported little overlap between HOXB13 and AR cistromes in PCa cell lines such as 22Rv1 and LN95 that express ARv7, an mCRPC-associated AR variant^19^. In these cells, HOXB13 protein interacts with ARv7 and they largely co-occupy the genome with ARv7, instead of full-length AR, to co-regulate target genes. HOXB13 co-localization with AR on the genome of PCa cells has also been shown enhanced upon the depletion of the chromodomain helicase DNA-binding protein 1 (CHD1)^20^. It is thus plausible that HOXB13 regulates AR cistrome in both a gene- and context-dependent manner. Likewise, the roles of HOXB13 in PCa tumorigenesis have also been an area of great controversy. Some studies have clearly shown that HOXB13 promotes cell growth^13, 16, 18, 19^, whereas others demonstrated growth-inhibitory effects^21, 22^. Norris et al. elegantly showed that HOXB13 knockdown (KD) abolished androgen-induced LNCaP cell growth but not that of hormone-deprived cells, suggesting context-dependent effects^16^. Further, germline mutation of HOXB13 at G84E has been associated with familial PCa and an early-onset disease^23^. Genome-wide association studies have identified PCa-linked SNPs locate within functional HOXB13 binding sites to disrupt HOXB13 regulation of target genes such as RFX6^13, 24–26^

Histone deacetylases (HDACs) are enzymes that catalyze the removal of acetyl groups from the lysine residues of histone proteins on nucleosomes, thus reducing histone acetylation and mediating transcriptional repression. HDAC3 is a member of the class I HDACs, which, also including HDAC1 and HDAC2, serve as the catalytic subunits of various co-repressor complexes^27^. HDAC3 is the catalytic subunit of the NCoR (Nuclear receptor corepressor)/SMRT (Silencing Mediator for Retinoid and Thyroid hormone receptors) complex^28^. The binding of NCoR and SMRT is, in turn, required to activate the histone deacetylase function of HDAC3^29–32^. These large multi-subunit protein complexes are recruited by tissue-specific transcription factors to selected enhancers and promoters to mediate transcriptional repression^33^. Recent studies have revealed a major function of HDAC3 in inhibiting *de novo* lipogenesis and controlling metabolic transcriptional networks in multiple tissue types^34, 35^. Inactivation of HDAC3 in the liver increased the expression of genes that drive lipid biosynthesis and storage, causing hepatomegaly and fatty liver^35, 36^. HDAC3 and NCoR1 co-localize near the regulatory elements of genes involved in lipid metabolism, suggesting a direct mechanism of transcriptional regulation^36^. A potential role of HDAC3-NCoR complex in regulating lipid metabolism in PCa cells, however, has not been previously explored.

Here we identify HDAC3 as an important cofactor of HOXB13 and they cooperate in remodeling the epigenome and suppressing *de novo* lipogenesis in PCa cells in both AR-positive and –negative cells. The MEIS domain of HOXB13 interacts with HDAC3, and this interaction is disrupted by HOXB13 G84E mutation, leading to restored expression of lipogenic regulators such as FASN in an AR-independent manner. Accordingly, HOXB13 loss results in massive lipid accumulation in PCa cells, thereby increasing cell motility *in vitro* and tumor metastasis *in vivo*. Lastly, we identify about 30% of mCRPC with low HOXB13 expression, which may be targeted by pharmaceutical inhibitors of FASN.

## RESULTS

### Genome-wide analysis revealed an essential molecular function of HOXB13 in suppressing *de novo* lipogenesis

To gain some insights into the genes and pathways that HOXB13 regulates in PCa cells, we generated stable LNCaP cell lines with control (pGIPZ), HOXB13 KD using shRNA targeting its 3’UTR (shHOXB13), and KD rescued by HOXB13 WT (shHOXB13+WT) or its G84E mutant (shHOXB13+G84E). RNA sequencing (RNA-seq) analysis of duplicate experiments identified 276 and 206 genes, with FDR<0.05 and fold change>=2.5, that were respectively decreased or increased by HOXB13 KD as compared to control samples (**Fig. 1A**). Re-introduction of WT HOXB13 to these cells fully rescued the gene expression changes. Interestingly, re-expression of HOXB13 G84E failed to rescue approximately one-third of the HOXB13-repressed genes, while its ability to rescue HOXB13-induced genes is mostly unimpaired. Gene Ontology (GO) analyses revealed that HOXB13-repressed genes were involved in steroid metabolic process, lipid biosynthetic process, and fatty acid metabolic process, whereas HOXB13-induced genes were enriched for molecular concepts related to cell proliferation, including cell division, cell cycle, and DNA replication (**Fig. 1B & S1A**). Further, RT-qPCR analyses of representative target genes confirmed that HOXB13 KD decreased the expression of master cell cycle regulators such as E2F1, FOXM1, MYBL2, and CDCA2^37^, which were fully rescued by the re-expression of either WT or G84E HOXB13 (**Fig. S1B**). By contrast, HOXB13 KD greatly increased the expression of lipogenic genes such as FASN, SREBF1/2, and PSA that has been shown to liquefy, in addition to semen, also lipids and citrate that fuel fatty acid synthesis^6^. Importantly, the expression of these HOXB13-repressed genes was abolished by re-introduction of WT, but not G84E, HOXB13 (**Fig. 1C**). Accordingly, western blotting (WB) confirmed that the protein levels of SREBF1, SREBF2, FASN, and PSA were also increased by HOXB13 KD, which was abolished by WT, but not G84E, HOXB13 re-expression (**Fig. 1D**). Taken together, our data indicate an important role of HOXB13 in suppressing de novo lipogenesis, a function that is impaired in the PCa-associated G84E mutant.

**Figure 1.**
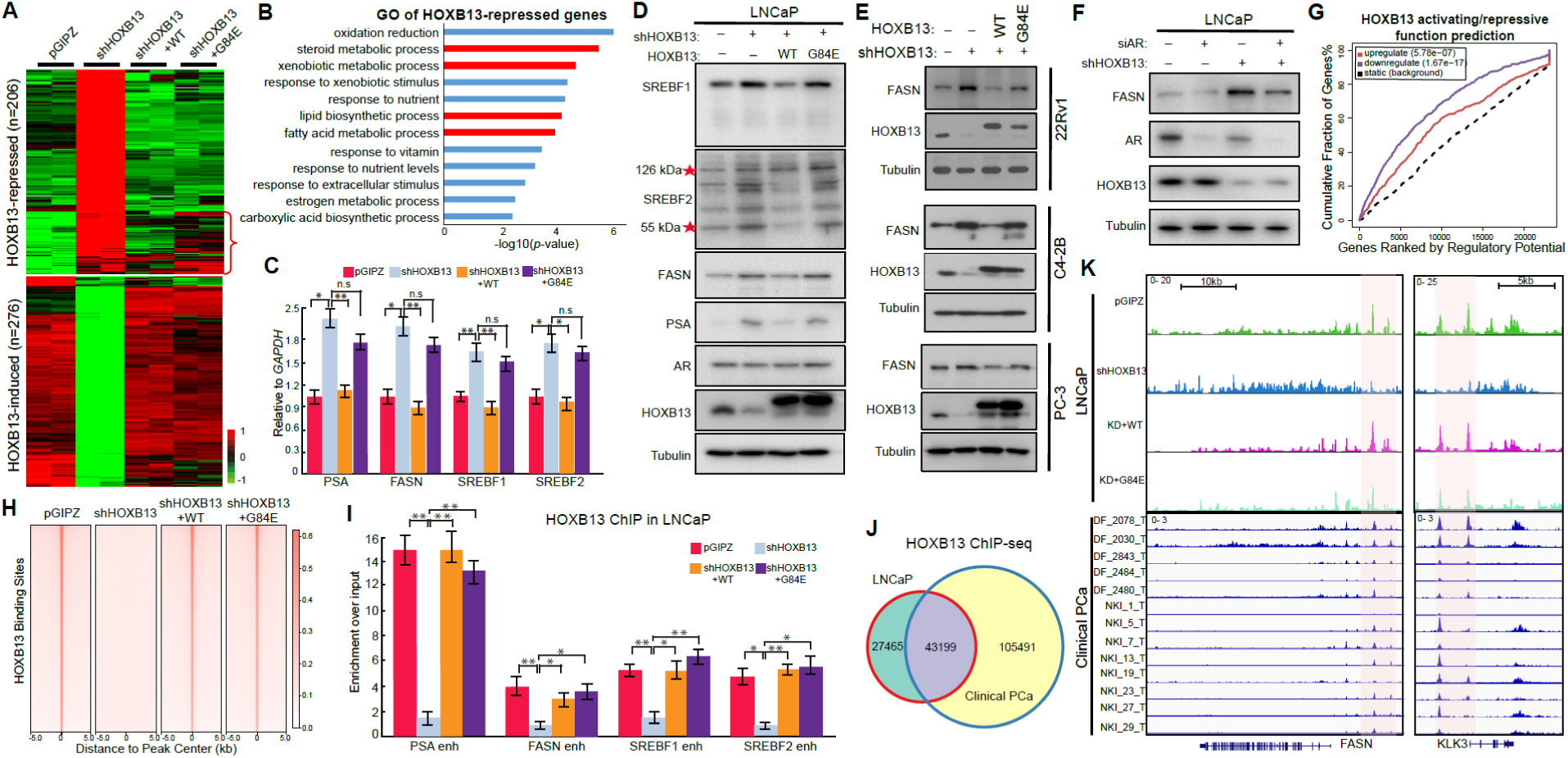
Genome-wide analysis revealed a major role of HOXB13 in suppressing lipid metabolism. **A.** Heatmap showing HOXB13-induced and -repressed gene expression in PCa cells with HOXB13 KD and/or rescue. LNCaP cells were infected with control pGIPZ, shRNA targeting HOXB13 (shHOXB13), shHOXB13 with concomitant infection by HOXB13-WT or G84E lentiviruses for 7 days. Cells were harvested for RNA-seq in replicate experiments. HOXB13-induced (n=277) and -repressed (n=207) genes were derived by comparing shHOXB13 with pGIPZ using FDR<0.05 and fold change>=2.5. Red bracket indicates HOXB13-repressed genes that are rescued by WT HOXB13, but not G84E mutant. **B.** GO analysis of HOXB13-repressed genes identified in (**A**). Top enriched molecular concepts are shown on Y-axis, while X-axis indicates enrichment significance. **C.** RT-QPCR validation of key lipogenic gene regulation by HOXB13 in LNCaP cells. Data were normalized to GAPDH. Data shown are mean (±SEM) of technical replicates from one representative experiment of three. Student’s t-test, \**p* < 0.05, \*\**p* < 0.01, n.s., not significant. **D.** WB of key lipogenic gene regulation by HOXB13 KD and rescue in LNCaP cells. *un-cleaved and cleaved forms of SREBF2. **E-F**. WB of FASN regulation by HOXB13 in multiple PCa cell lines with AR and/or HOXB13 KD. **G.** HOXB13 ChIP-seq data was integrated with RNA-seq profiling of control and HOXB13 KD LNCaP cells using BETA software to predict activating/repressive function of HOXB13. Genes are cumulated by the rank on the basis of the regulatory potential score from high to low. The dashed line indicates the nondifferentially expressed genes as background. The red and the purple lines represent the upregulated and downregulated genes, respectively, and their P values were calculated comparing with the background group by the Kolmogorov-Smirnov test. **H.** Heatmap showing HOXB13 ChIP-seq intensity in HOXB13 KD and rescue LNCaP cells. **I.** HOXB13 ChIP-qPCR of lipogenic genes in cells with HOXB13 de-regulation. **J.** Venn Diagram comparing HOXB13 cistromes between LNCaP and human PCa tissues **K.** Genome browser track showing HOXB13 occupancy at lipogenic genes in LNCaP and clinical PCa.

To determine whether HOXB13-regulated pathways are dependent on AR, we analyzed hormone-deprived LNCaP cells with control and HOXB13 KD. Interestingly, GO analysis demonstrated that, under androgen-depleted condition, i.e., without AR activation, HOXB13-repressed genes remained to be strongly enriched in lipid and fatty acid metabolism (**Fig. S1C**), whereas HOXB13-induced genes were no long enriched for cell proliferation (**Fig. S1D**). To further investigate the differential dependence of HOXB13-induced and –repressed transcriptional programs on AR signaling, we performed RNA-seq analysis of the AR-negative but HOXB13-positive PC-3 cells. GO analysis demonstrated that HOXB13-repressed genes in PC-3 cells continued to strongly enrich for lipid and fatty acid metabolism, whereas HOXB13-induced genes were not enriched for cell cycle/growth concepts (**Fig. S1E-F**). To confirm the newly identified, AR-independent role of HOXB13 in regulating lipid metabolism, we examined additional cell lines including PC-3 and CRPC cell lines C4-2B and 22Rv1. Our data validated that HOXB13 KD in these cells increased FASN expression, which was abolished by re-expression of WT, but not G84E, HOXB13 (**Fig. 1E**). Moreover, HOXB13 depletion continued to up-regulate FASN in LNCaP cells with AR knockdown, supporting an AR-independent mechanism (**Fig. 1F**, lanes 2 and 4).

Lastly, to determine whether HOXB13 directly controls these lipogenic regulators, we performed HOXB13 ChIP in LNCaP cells. ChIP-seq demonstrated that a majority of HOXB13 binding sites are at enhancer regions within introns (46% peaks) or between genes (47%) (**Fig. S1G**). Integration of ChIP-seq and transcriptomic data using BETA software^38^ predicted that transcriptional repression is the prominent molecular function of HOXB13, much more than its ability in gene activation (**Fig. 1G**). Further, ChIP-seq in LNCaP cells with HOXB13 deregulation showed that HOXB13 binding sites were drastically reduced in shHOXB13 cells, as expected, and largely rescued by re-expression WT or G84E HOXB13 (**Fig. 1H**). ChIP-qPCR using gene-specific primers confirmed that HOXB13 bound to the regulatory elements of PSA, FASN, SREBF1, and SREBF2, which were abolished by HOXB13 KD (**Fig. 1I**), supporting that they are direct targets of HOXB13. Importantly, re-introduction of either WT or G84E HOXB13 rescued HOXB13 occupancy at these genes, indicating that G84E mutation did not affect the DNA-binding ability of HOXB13, which is consistent with prior reports that HOXB13 interacts with DNA through its HOX DNA binding domain^16^. It also suggests that the impaired ability of G84E HOXB13 to repress the expression of these genes **(Fig. 1C**) may be mediated by cofactors rather than HOXB13 itself. Being consistent with the ability of HOXB13 to repress lipid metabolism in an AR-independent manner, we found that HOXB13 also bound to the enhancers of lipogenic regulators in PC-3 cells (**Fig. S1H**). Importantly, comparison of HOXB13 binding sites with those identified in clinical specimens ^39^ revealed significant overlap and confirmed HOXB13 occupancy at the enhancers of lipogenic genes in human PCa tissues (**Fig.1J-K**).

Taken together, our data identified an essential role of HOXB13 in directly binding to lipogenic enhancers to suppress *de novo* lipogenesis in PCa cells - a function that is independent of AR but may be associated with essential cofactors that are disrupted by G84E mutation.

### The MEIS domain of HOXB13 protein interacts with HDAC3 protein

To identify potential cofactors of HOXB13 in suppressing lipogenic programs, we performed tandem affinity purification followed by mass spectrometry analysis of WT and G84E HOXB13 expressed in LNCaP cells. Out of the HOXB13-enriched proteins are previously reported interactors such as AR and its cofactors FOXA1, GATA2, and NKX3.1 (**Fig. S2A**). However, these interactions were not strongly disrupted by G84E as compared to WT HOXB13. Interestingly, we found strong interactions of WT HOXB13 with HDAC1/3 and their corepressors NCoR1/2 and TBL1X. Importantly, these interactions were all drastically reduced by G84E mutation as supported by peptide counts **(Fig. 2A**). To confirm this, we performed co-immunoprecipitation (co-IP) of Flag-tagged HDAC1 or HDAC3 in LNCaP cells and confirmed that HOXB13 interacts with both of them (**Fig. 2B**). In a reciprocal experiment, co-IP experiment showed that HA-WT HOXB13 strongly interacted with both HDAC1 and HDAC3, which, especially HDAC3, barely interacts with HA-G84E HOXB13 (**Fig. 2C**). In contrast, G84E mutation did not affect the ability of HOXB13 to interact with AR.

**Figure 2.**
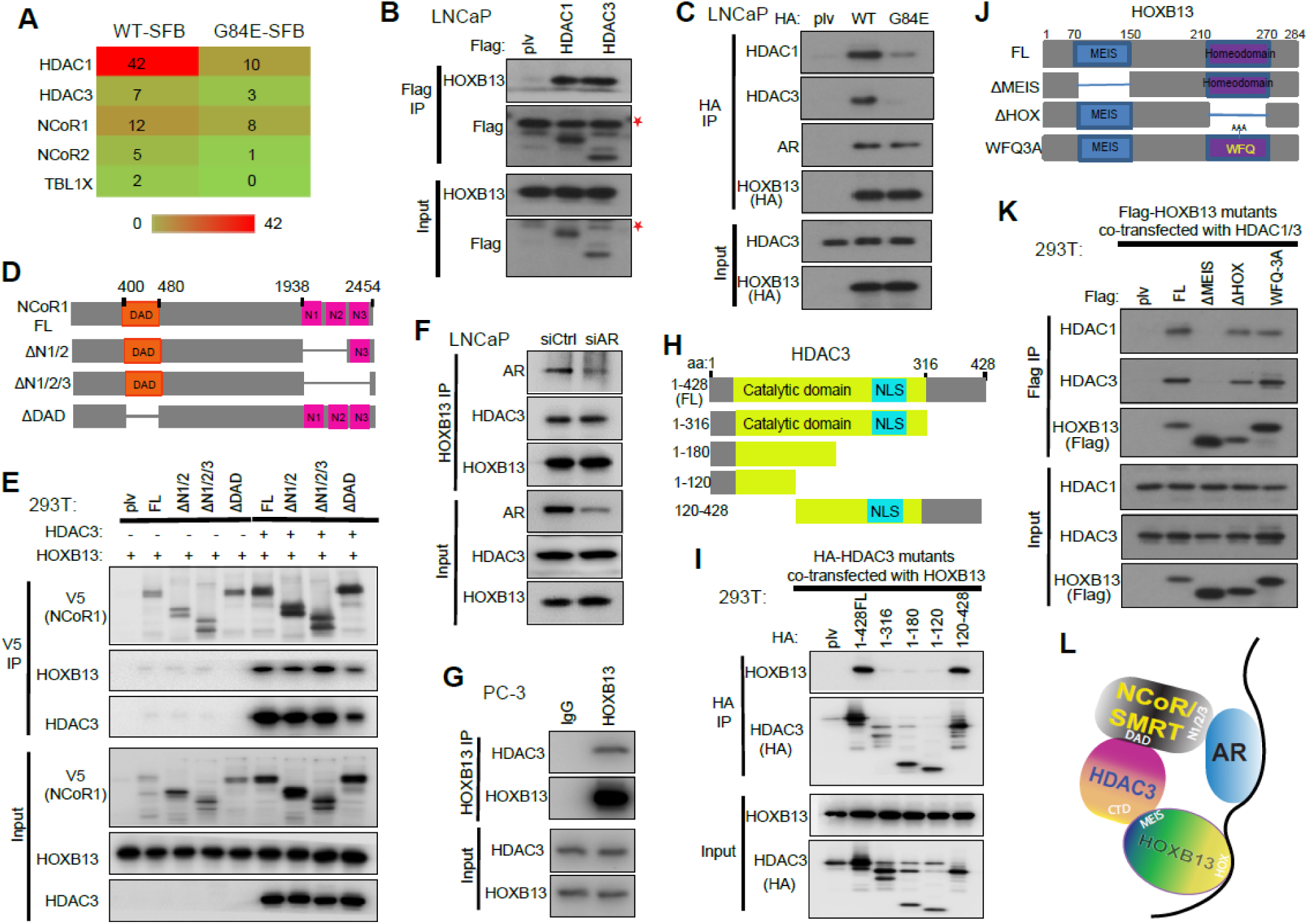
The MEIS domain of HOXB13 protein interacts with HDAC3 protein. **A.** Heatmap showing interactor proteins of HOXB13 WT or G84E mutant. Whole cell lysates of LNCaP cells stalely expressing SFB-tagged HOXB13-WT or G84E mutant were subjected to two rounds of sequential purification and then subjected to mass spectrometry analysis. **B.** Co-IP of HDAC1 and 3 shows interaction with HOXB13. Whole cell lysates from LNCaP cells stably expressing Flag-tagged HDAC1 or HDAC3 were subjected to co-IP using anti-Flag antibody. The eluted co-IP complex and input were analyzed by WB. Asterisk indicates a non-specific band detected by anti-Flag antibody. **C.** Whole cell lysates (input) from LNCaP stably expressing HA-HOXB13-WT or -G84E were subjected to co-IP using an anti-HA antibody. **D-E**. Co-IP of V5-NCoR1 FL or deletion mutants (**D**) expressed in 293T cells along with HOXB13, with or without HDAC3 co-expression (**E**). **F.** HOXB13 interaction with HDAC3 is NOT dependent on AR. Whole cell lysates from LNCaP with control (siCtrl) or AR knockdown (siAR) were subjected to co-IP using anti-HOXB13 antibody. **G.** Co-IP of HOXB13 in the AR-negative PC-3 cells. **H-I**. Co-IP of HA-HDAC3 FL or deletion mutants (**H**) in 293T cells with co-expression of HOXB13 (**I**). **J-K**. Co-IP of Flag-HOXB13 FL or deletion mutants (**J**) in 293T cells with co-expression of HDAC1 and HDAC3 (**K**). **L**. A model depicting the interactions between HOXB13, AR, and HDAC3/NCoR complex.

HDAC3 is the catalytic subunit of the NCoR/SMRT complex, which interacts with lineage transcription factors such as AR through its N-terminal nuclear receptor interaction domain^30, 31, 40, 41^. To investigate whether HDAC3 interaction with HOXB13 is also mediated by NCoR/SMRT, we overexpressed NCoR1 and its various deletion mutants (**Fig. 2D**) in 293T cells along with co-expression of HOXB13 and/or HDAC3. Surprisingly, co-IP showed very weak interactions between HOXB13 and NCoR1 in cells without HDAC3 co-expression, suggesting that HOXB13 and NCoR1 do not directly interact (**Fig. 2E**). In cells with concurrent expression of ectopic HDAC3, NCoR1 IP expectedly pull down HDAC3, which was markedly decreased by the deletion of DAD domain of NCoR1 (ΔDAD), a domain previously reported to mediate NCoR1/SMRT interaction with HDAC3^42, 43^. Importantly, in these cells a strong interaction between HOXB13 and NCoR1 was detected and their interaction was also mitigated by ΔDAD. This evidence support that HOXB13 interacts with HDAC3 first, which subsequently recruits NCoR1, whose binding is critical for activating the enzymatic activities of HDAC3^31^. This data reveals HOXB13-HDAC3-NCoR1 as a complex that is bridged by HDAC3, being very different from the NCoR-mediated HDAC3-NCoR-AR complex. Further, co-IP in LNCaP cells with AR KD showed that AR KD did not affect the interaction between endogenous HOXB13 and HDAC3 proteins (**Fig. 2F**). HOXB13 and HDAC3 protein interaction was also detected in the AR-negative PC-3 cells, suggesting its independence in AR (**Fig. 2G**).

We next sought to determine which domains of HDAC3 interact with HOXB13. We cloned HDAC3 full-length (FL) and a series of truncation mutants (aa1-316, aa1-180, aa1-120 and aa120-428) (**Fig. 2H**), which was co-transfected with HOXB13 in 293T cells. Co-IP revealed strong enrichment of HOXB13 with FL (aa1-428) and aa120-428 fragment of HDAC3, but not its N-terminal fragments, suggesting that the C-terminal domain (CTD) of HDAC3 is required for its interaction with HOXB13 (**Fig. 2I**). Conversely, HOXB13 proteins contain two important functional domains: C-terminal HOX DNA binding domain (aa217-275) and N-terminal MEIS domain (aa80-91 and aa136-146) that is responsible for its interaction with MEIS proteins^22^. To examine how these functional domains affect HOXB13 interaction with HDAC3, we generated several deletion mutants (**Fig.2J**), including ΔMEIS (aa70-150), ΔHOX (aa210-270) and HOXB13^WFQ > AAA^ mutation (WFQ-3A) that were previously shown to disrupt HOXB13 DNA binding ^16^ (**Fig. S2B**). Our data showed that ΔMEIS, but not its DNA-binding mutants, abolished HOXB13 interaction with HDAC3 (**Fig. 2K**). Similarly, co-IP experiments of a series of N-terminal truncation mutants of HOXB13 revealed strong HDAC3 interactions with HOXB13 FL and aa70-284 fragment, but only weak interaction with aa150-280 and aa210-284 (**Fig. S2C-D**). These results are consistent with our earlier finding that G84E mutant, located within the MEIS domain, has impaired ability to interact with HDAC3. In summary, our data suggest a model wherein HOXB13 MEIS domain interacts with the CTD of HDAC3, which subsequently recruit NCoR1 for enzymatic activation, and that this complex is independent of AR (**Fig. 2L**).

### HOXB13 recruits HDAC3 to target chromatin to catalyze histone deacetylation

To examine the effects of HOXB13 and HDAC3 protein interaction on chromatin, we performed ChIP-seq in LNCaP cells and indeed observed substantial overlap between HOXB13 and HDAC3 cistromes, a majority of which are distinct from HOXB13 and AR co-occupied sites (**Fig.3A**). By contrast, the overlap between HOXB13 and HDAC1 binding sites was much limited (**Fig. S3A**). This could partially be explained by preferential binding of HDAC3, similar as HOXB13, at intronic and intergenic enhancers (**Fig. S3B**), while HDAC1 was previously shown to occupy more at gene promoters ^44^. Further, HOXB13 and HDAC3 co-occupied sites, defined hereafter as HH sites, are enriched for H3K27ac and increased chromatin accessibility as determined by Assay for Transposase-Accessible Chromatin using sequencing (ATAC-seq) (**Fig.3B**), being consistent with previous reports that HDAC3 is targeted to active genes with acetylated histone^45^. Critically, depletion of HOXB13 abolished HOXB13 cistrome, clearly reduced HDAC3 recruitment to the chromatin, and increased H3K27ac, while AR binding, in contrast, was not altered (**Fig. 3B & S3C**). Analyses of H3K27ac ChIP-seq performed in clinical PCa specimens^39^ confirmed strong H3K27ac at HH sites in human PCa, supporting their clinical relevance (**Fig.3C**). We thus hypothesized that HOXB13 recruits HDAC3 to active chromatin for histone deacetylation and transcriptional repression. Indeed, there was a global reduction of HDAC3 binding sites in HOXB13-depleted cells and the lost sites were more strongly enriched for HOXB13 co-occupancy (**Fig. 3D**).

**Figure 3.**
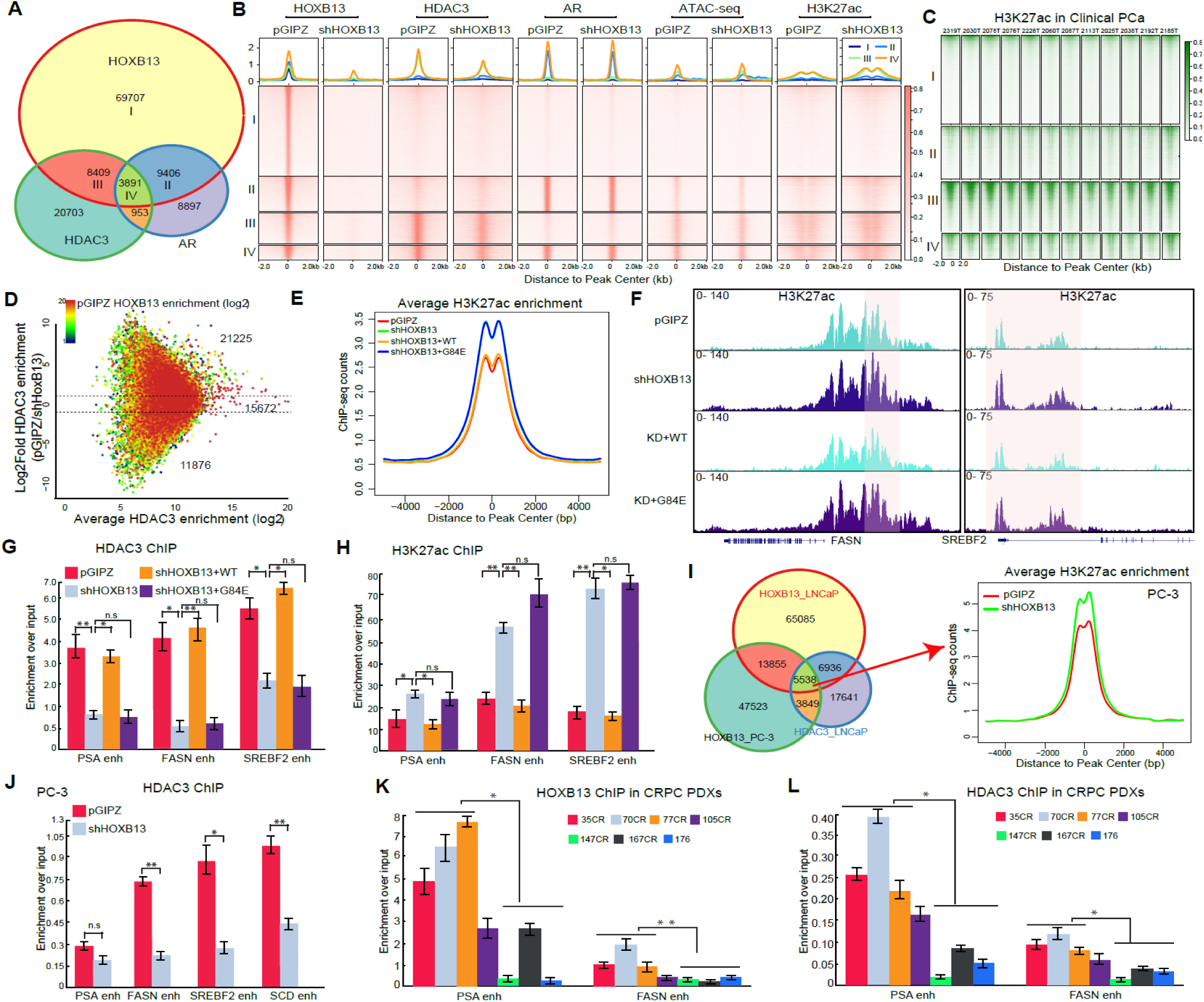
HOXB13 recruits HDAC3 to target chromatin to catalyze histone deacetylation. **A**. Venn diagram showing overlap of HOXB13, AR and HDAC3 binding sites in LNCaP cells. **B-C**. Heatmap showing signals of HOXB13, HDAC3, AR, and H3K27ac ChIP-seq and ATAC-seq in LNCaP cells with control or HOXB13 KD (**B**) and of H3K27ac ChIP-seq in a set of 12 clinical PCa samples (GSE130408), centered around (±2kb) selected genomic regions defined in **A**. Color bar on the right shows the scale of enrichment intensity. **D.** MA plot showing differential HDAC3 binding sites in control (pGIPZ) and HOXB13 KD (shHOXB13) LNCaP cells. Color encodes the intensity of HOXB13 ChIP-seq in control cells. Dotted lines represent 2-fold differences. **E.** Average H3K27ac signal centered around (±5 kb) HDAC3 and HOXB13 co-occupied sites. H3K27ac ChIP-seq were performed in LNCaP infected with pGIPZ, shHOXB13 (KD), or KD with co-infection of WT or G84E HOXB13. **F.** Genome browser view of H3K27ac ChIP-seq signal around representative lipogenic genes. **G-H.** HDAC3 (**G**) and H3K27ac (**H**) ChIP-qPCR validation of lipogenic gene enhancers (enh) in LNCaP with HOXB13 KD and/or rescue. Data were normalized to 2% of input DNA. Shown are mean (±SEM) of technical replicates from one representative experiment of three. Student’s t-test, \**p* < 0.05, \*\**p* < 0.01, *n.s.* not significant. **I.** Venn diagram showing overlapping HDAC3 and HOXB13 cistromes in PC-3 and LNCaP cells (left). H3K27ac ChIP-seq was performed in PC-3 cells with control or HOXB13 KD and their average intensity plots centered around (±5 kb) co-occupied sites are shown on the right. **J.** HDAC3 ChIP-qPCR of lipogenic gene enhancers in PC-3 cells with control or HOXB13 KD. Data were processed and plotted as in **G-H**. **K-L**. HOXB13 (**K**) and HDAC3 (**L**) ChIP-qPCR analysis of lipogenic enhancers in mCRPC LuCaP PDXs. Data were processed and plotted as in **G-H**.

To examine how HOXB13 WT and G84E mutants differentially modulate histone de-acetylation, we performed H3K27ac ChIP-seq in LNCaP cells with HOXB13 KD and/or rescue by WT or G84E. Bioinformatics data analyses of HH sites revealed markedly increased H3K27ac following HOXB13 KD, which was again decreased by WT re-expression (**Fig. 3E-F**). However, HOXB13 G84E exhibited an impaired ability to suppress H3K27ac. To confirm this, we performed ChIP-qPCR analysis using primers specific to lipogenic enhancers. The results indicated that HDAC3 recruitment to these genes was decreased upon HOXB13 KD, which was rescued by re-expression of WT, but not G84E HOXB13 (**Fig. 3G**). Accordingly, re-expression of WT HOXB13, but not G84E, abolished H3K27ac (**Fig. 3H).** To demonstrate the molecular functions of HOXB13 in an AR-independent context, we performed HOXB13 ChIP-seq in the AR-negative PC-3 cells. We observed over 70,000 HOXB13 binding sites in PC-3 cells, 27.4% of which were also bound HOXB13 in LNCaP cells (**Fig. 3I**). Importantly, around 44% of HH sites found in LNCaP cells were also bound by HOXB13 in PC-3 cells and showed increased H3K27ac upon HOXB13 depletion. ChIP-qPCR validated HDAC3 occupancy at the regulatory elements of lipogenic genes in PC-3 cells, which was abolished by HOXB13 KD similar as in LNCaP cells (**Fig. 3J**). Accordingly, H3K27ac at these genes were increased in HOXB13-KD cells (**Fig. S3D**). To validate the HOXB13/HDAC3-H3K27ac axis in human PCa cells, we performed ChIP analyses of tumor bits from 7 Patient-Derived Xenografts (PDXs) established from human metastatic castration-resistant PCa (mCRPC)^46^ that can be separately into 3 high-, 3 low-, and 1 intermediate-HOXB13 levels (**Fig. S3E-F**). ChIP-qPCR analyses confirmed significantly higher HOXB13 and HDAC3 occupancy at the lipogenic gene enhancers in HOXB13^high^ than HOXB13^low^ tumors (**Fig. 3K-L**). Conversely, H3K27ac at these sites was stronger in HOXB13^low^ tumors (**Fig. S3G**). Altogether, our results demonstrated that HOXB13 recruits HDAC3 to lipogenic enhancers to catalyze histone deacetylation of target genes.

### HDAC3 is required for HOXB13-mediated suppression of *de novo* lipogenesis

To examine whether HDAC3, like HOXB13, regulates lipid/fatty acid metabolisms in PCa cells, we performed HDAC1/3 KD in LNCaP cells and found HDAC3 as a main regulator of FASN and PSA, albeit depletion of HDAC1 also slightly increased their expression (**Fig. S4A**). RNA-seq analyses showed that HDAC1/3 KD significantly altered the expression of 683 genes, which overlapped with 36% of HOXB13-regulated genes, and they were either co-repressed or co-induced (**Fig. 4A & S4B**). In agreement with this, GSEA analysis of HDAC1/3-repressed genes showed highly significant enrichment for up-regulation in shHOXB13 cells, indicating their being co-repressed by HOXB13 (**Fig. 4B**). GO analyses of HDAC1/3-target genes demonstrated enrichment of molecular concepts that were very similar to HOXB13-target genes. Namely, HDAC1/3-repressed genes were involved in metabolic pathways such as oxidation reduction, fatty acid metabolic process, and steroid metabolic process, while HDAC1/3-induced genes are strongly enriched in cell division and mitotic processes **(Fig. 4C & S4C**). This data suggests that HDAC3 is likely an important mediator of HOXB13 function.

**Figure 4.**
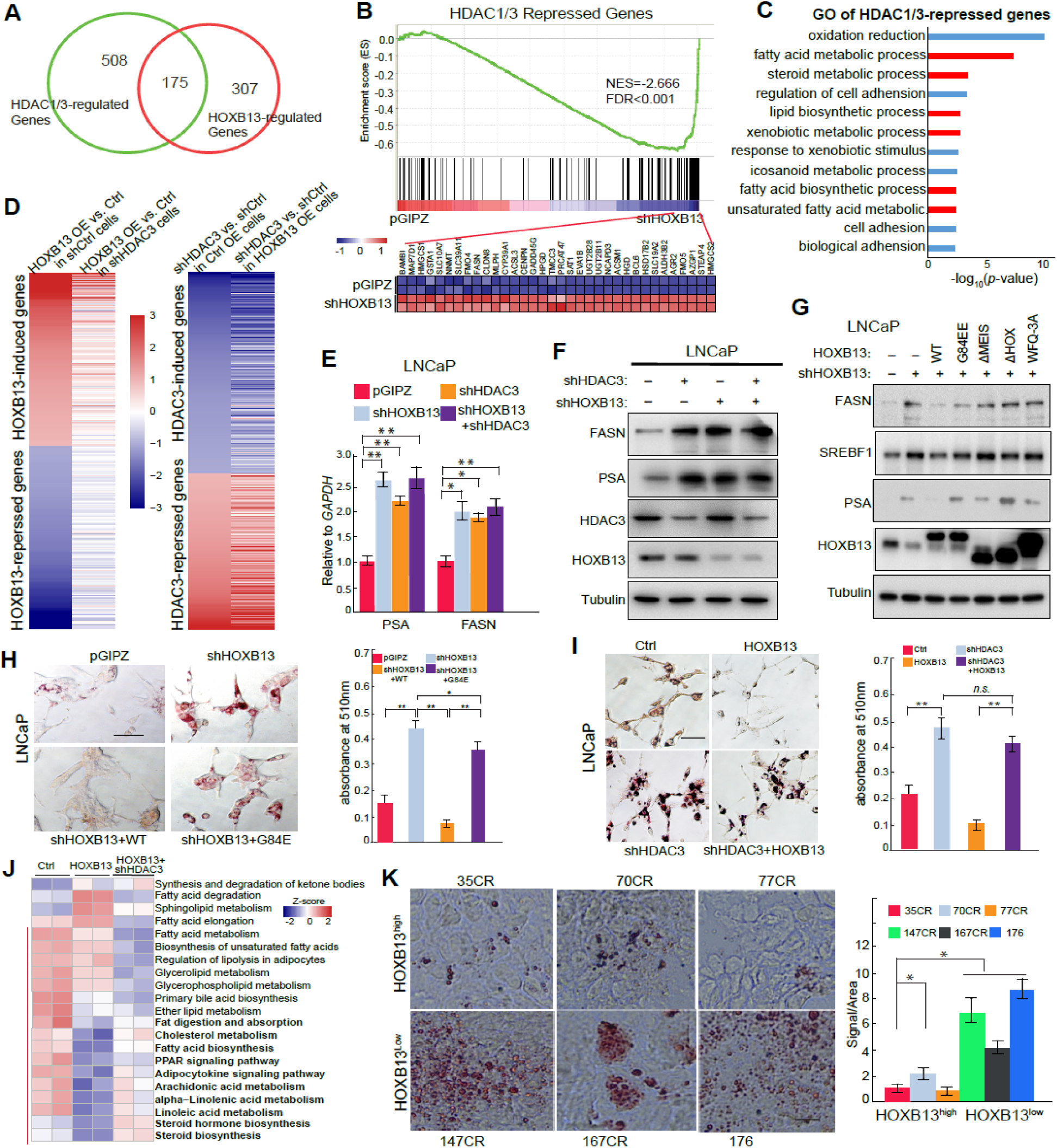
HDAC3 is required for HOXB13-mediated suppression of *de novo* lipogenesis. **A.** RNA-seq was performed in LNCaP cells with control, HOXB13 KD, or HDAC1/HDAC3 double KD. Differentially regulated genes were identified by fold change >=2 and FDR<0.05. **B.** GSEA showing the enriched expression of HDAC1/3-repressed genes in pGIPZ LNCaP cells. Shown at the bottom are heatmaps of leading-edge gene expression in corresponding samples. **C.** GO analysis of HDAC1/3-repressed genes was performed using DAVID. Top enriched molecular concepts are shown. X-axis indicates enrichment significance. **D.** HOXB13 regulation of target genes is dependent on HDAC3, but not vice versa. Heatmaps shown on the left indicates HOXB13 regulation of its induced (top) and repressed (bottom) genes in shCtrl or shHDAC3 LNCaP cells. Heatmaps on the right show HDAC3 regulation of its induced (top) and repressed (bottom) genes in LNCaP cells with control or HOXB13 OE. **E-F**. RT-QPCR and WB analysis of FASN and PSA expression in LNCaP cells with KD of HOXB13, HDAC3, or both. RT-QPCR data were normalized to GAPDH. Data shown are mean (±SEM) of technical replicates from one representative experiment of three. Student’s t-test, \**p* < 0.05, \*\**p* < 0.01, *n.s.* not significant. **G.** WB analysis of HOXB13 and HDAC1/3 co-regulated genes in LNCaP cells with HOXB13 KD and rescue by HOXB13 WT or mutants. Tubulin was used a loading control. **H.** Representative images of Oil Red O staining (left) and quantification (right) of neutral lipids in LNCaP with shHOXB13 and rescue by WT or G84E HOXB13. *Scale bar*: 50 µm. Data shown are mean (±SEM) of triplicate wells. *\*\*p*<0.01, \**p*<0.05, *n.s.* not significant by Student’s *t*-test. **I.** Oil Red O staining (left) and quantification (right) of neutral lipids in LNCaP cells with control, HOXB13 OE and/or HDAC3 KD. Data analyzed and presented as in **H**. **J.** HOXB13 or HDAC3 regulation of KEGG lipid metabolism-related pathways. A total of 21 lipid metabolism-related gene sets were downloaded from KEGG. For each gene set, a subset of genes that are differentially regulated (FDR <0.05, fold>2) by HOXB13 were selected and Z-score for each gene normalized across all 6 samples. Heatmap shows the average Z-score of each gene set (row) in each sample (column). Red bar on the left indicates lipid metabolism pathways that are repressed by HOXB13 OE. Concepts that were rescued by shHDAC3 are shown in bold (right). **K.** Oil Red O staining (left) of neutral lipids in HOXB13^low^ or HOXB13^high^ LuCaP PDXs. Data was quantified (right) using Image J as percentage staining per area. Data analyzed as in **H**.

Next, we sought to determine whether HDAC3, through chromatin remodeling, is required for HOXB13-mediated regulation of downstream genes. To this end, we analyzed RNA-seq data comparing control and HOXB13 overexpression (OE) in HDAC3-expressing or - depleted LNCaP cells. We identified 162 and 172 genes that were respectively induced and repressed by HOXB13 in the control, HDAC3-expressing, cells (**Fig. 4D**). Analysis of the same gene set in HDAC3-depleted cells revealed significantly reduced differential expression by HOXB13 OE, suggesting a dependency on HDAC3. Further, HDAC3 KD increased and decreased a respective set of 386 and 297 genes in control cells, many of which remained sensitive to HDAC3 in HOXB13-OE cells, supporting HDAC3 as a direct regulator of gene expression. Conversely, to determine whether HOXB13 is required for the activity of HDAC3 on gene regulation, we performed KD of HOXB13, HDAC3, or both in LNCaP cells. RT-QPCR and WB analyses revealed that depletion of either HOXB13 or HDAC3 alone was sufficient to restore PSA and FASN expression and knockdown of both genes did not further increase target gene expression (**Fig. 4E-F**). In the absence of HOXB13, HDAC3 failed to regulate lipogenic genes, and vice versa, suggesting that both HOXB13 and HDAC3 are required for transcriptional repression of lipogenic genes. This supports our model showing the necessity of HOXB13 to recruit HDAC3 to the enhancers and of HDAC3 to catalyze histone de-acetylation for gene repression. The same trend of regulation was also observed in the AR-negative PC-3 cells, supporting this as an AR-independent function (**Fig. S4D**). To further confirm this, we performed HOXB13 KD in LNCaP cells followed by rescue using various mutants. As controls, HOXB13 depletion expectedly led to up-regulation of FASN, SREBF1, and PSA, which were again repressed by re-introduction of WT HOXB13 (**Fig. 4G**). HOXB13 ΔHOX/WFQ-3A mutants, although capable of interacting with HDAC3, are unable to bind DNA^16^ and consequently failed to repress lipogenic genes. Importantly, in contrast to WT HOXB13, ΔMEIS and G84E mutants, which have impaired ability to interact with HDAC3, also failed to fully repress these genes. Together, our data support a model wherein HOXB13 interacts with and recruits HDAC3 to target chromatin for transcriptional repression.

To elucidate the roles of HOXB13 in regulating lipid phenotypes in PCa, we performed untargeted lipidomic analysis by Quadrupole Time of Flight (Q-TOF) Liquid Chromatography (LC)/Mass Spectrometry (MS) of LNCaP cells with control, shHOXB13, KD with WT or G84E re-expression. The results revealed that most of the lipids were increased following HOXB13 KD and rescued by the re-expression of WT, but not G84E mutant, HOXB13 (**Fig. S4E**). Among them, glycerophospholipid (PG) is a key component of biological membranes that has been shown to be the most significantly upregulated lipid species in MYC-driven cancer ^47^, while Cholesterol and Triglycerides (TG) are main components of lipid droplets and have been reported up-regulated in aggressive PCa^7^. Similarly, HOXB13 WT, but not G84E, was found to repress free fatty acids such as oleic acid and linoleic acid (**Fig. S4F**). Next, we sought to investigate whether such regulation of lipid metabolism by HOXB13 leads to changes in lipid accumulation, which has been observed in aggressive PCa cells^3, 7, 8^. Oil Red O (ORO) staining of neutral triglycerides and lipids detected greatly increased lipid accumulation in HOXB13-depleted LNCaP cells, which was fully abolished by re-expressing WT HOXB13, but only partially rescued by G84E mutant (**Fig. 4H**). ORO staining of additional CRPC cell line 22Rv1 and AR-negative cell line PC-3 revealed similar mode of regulation, supporting that HOXB13 suppresses lipid accumulation independently of AR and that this functionality is disrupted by G84E mutation (**Fig. S4G-H**). To determine whether this function is mediated by HDAC3, we performed ORO staining of LNCaP cells following HOXB13 OE in control or HDAC3-depeleted cells (**Fig. 4I**). Our results demonstrated that HOXB13 OE markedly reduced lipid accumulation in the control cells, but had very limited effect in cells with concurrent depletion of HDAC3. Collectively, these data support that HOXB13 suppresses lipid accumulation through a mechanism that is independent of AR but requiring HDAC3.

As lipid metabolism involves many complex molecular processes, we examined 21 lipid metabolism-related pathways denoted in the Kyoto Encyclopedia of Genes and Genomes (KEGG) database. To investigate how each metabolic pathway is regulated by HOXB13 and/or HDAC3, we computed a metabolic pathway score for each of the 21 lipid metabolism pathways based on the average Z-scores of expression of all genes within each pathway in LNCaP cells with control, HOXB13 OE, or OE with concomitant shHDAC3. Clustering analysis of the metabolic pathway scores revealed that metabolic pathways associated with fatty acid biosynthesis, fat digestion and absorption, steroid biosynthesis were repressed by HOXB13 OE and their expression was restored upon concurrent HDAC3 KD (**Fig. 4J**). Collecting genes from these molecular concepts, we generated a lipogenic gene signature. GSEA analysis of this lipogenic gene signature confirmed their strong enrichment for repression by HOXB13 OE and rescue by HDAC3 depletion (**Fig. S4I**). Lastly, ORO staining of human PDX tumors confirmed much stronger lipid accumulation in HOXB13^low^ than HOXB13^high^ PCa (**Fig. 4K**). Taken together, our data indicate that HOXB13 and HDAC3 cooperatively suppress *de novo* lipogenesis and reduce lipid accumulation in PCa cells.

### HOXB13 is hypermethylated and down-regulated in CRPC

To investigate the clinical relevance of this pathway in human PCa, we examined HOXB13 expression in publically available PCa expression profiling datasets and observed significant decreases of HOXB13 mRNA in metastatic CRPC relative to localized PCa in multiple large datasets (**Fig. 5A**). Of noteworthy, we observed a transient up-regulation of HOXB13 during initial PCa development, as compared to benign prostate, which is consistent with its reported role as an essential cofactor of AR, a key driver of PCa growth. To confirm these expression patterns at protein levels, we performed immunohistochemical (IHC) staining of HOXB13 using tissue microarray (TMA) of human PCa tissues (**Fig.5B-C**). Consistent with its role as a transcription factor, HOXB13 showed strong and punctuated nuclear staining. More than 95% of primary PCa showed moderate to strong staining for HOXB13, while only 33% of metastatic CRPC tumors exhibited similar levels of high HOXB13 (**Fig. 5C**), suggesting HOXB13 loss or down-regulation in a significant subset of mCRPC tumors.

**Figure 5.**
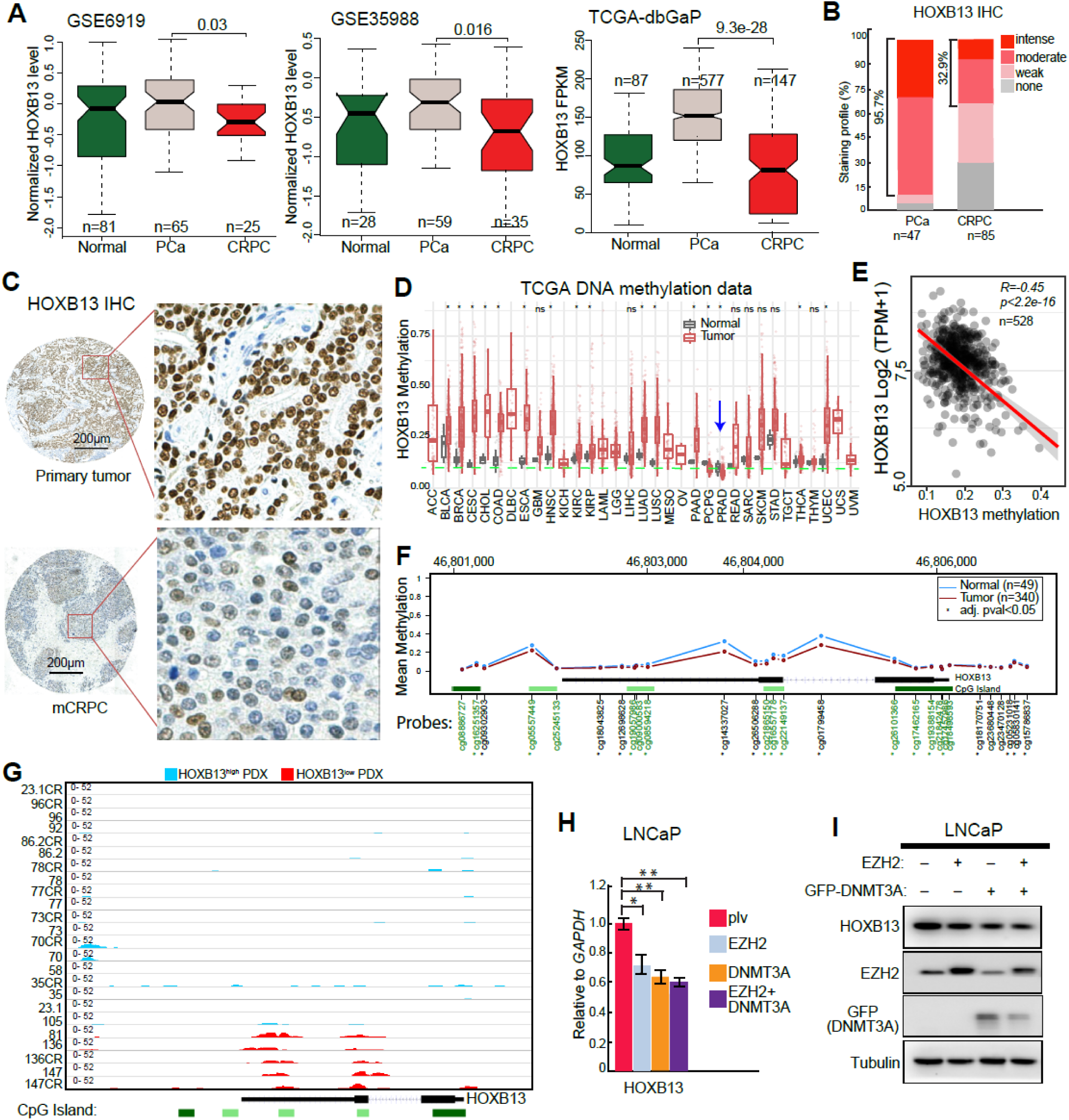
HOXB13 is hypermethylated and down-regulated in CRPC. **A.** HOXB13 expression levels in normal prostate, primary PCa, and metastatic CRPC in publically available PCa expression profiling datasets. *P* values between primary and metastatic PCa were calculated using Student’s t test. **B.** Quantification of HOXB13 IHC staining intensities in human PCa and CRPC. Y axis shows percentage of tumors with none, weak, moderate, and strong IHC staining. **C.** Representative images of IHC analyses of human primary PCa and mCRPC tissues. Representative images of HOXB13 staining in primary tumors (top panels) and metastatic CRPC (bottom panels) are shown in low- (×40) and high-magnification (×200). **D.** HOXB13 gene methylation levels in a variety of cancer types and corresponding benign samples using TCGA data. Blue arrow indicates prostate cancer (PRAD). *marks tumor types of significant differences with matched benign samples. n.s. not significant. **E.** Correlation between HOXB13 gene methylation (X-axis) and expression at mRNA level, indicated by transcripts per million (TPM) on Y-axis. **F.** Average methylation levels of Illumina Infinium probes targeting different regions of the HOXB13 gene. Probes in green fonts target CpG islands. *probes with significantly differential methylation between normal and prostate tumor samples. **G.** Genome Browser track showing HOXB13 gene methylation in CRPC tumors. LuCaP PDX tumors with high (blue) and low (red) HOXB13 expression levels were analyzed by MeDIP-seq. **H-I**. RT-QPCR and WB analyses of LNCaP cells subjected to EZH2 and GFP-DNMT3A overexpression. PCR data shown are mean (±SEM) of technical replicates from a representative experiment of three. Student’s t-test, *p < 0.05, **p < 0.01, n.s. not significant.

To gain some insights of potential mechanisms of HOXB13 loss of expression, we analyzed genome-wide DNA methylation data generated by The Cancer Genome Atlas (TCGA) program using Illumina Infinium Human Methylation 450K BeadChip. Utilizing the SMART App^48^, comparison across 23 normal tissue types revealed markedly less methylation of the HOXB13 gene in the prostate tissue **(Fig. 5D**), being consistent with previous reports of exclusive HOXB13 expression in the prostate^14, 15^. Interestingly, HOXB13 methylation is further decreased in primary PCa (PRAD) as compared to normal, which is in agreement with our earlier observation of relatively higher HOXB13 mRNA expression in the former (**Fig.5A**). Indeed, HOXB13 expression is significantly, and negatively, associated with HOXB13 methylation (**Fig.5E**), supporting methylation as an important mechanism to regulate HOXB13 gene expression. Analysis of TCGA-PRAD methylation data using Wanderer tool^49^ revealed three methylation hotspots on the HOXB13 gene, located near CpG islands, that showed significantly decreased methylation in prostate tumors as compared to normal (**Fig.5F**).

To gain some insights to the levels of HOXB13 methylation in CRPC tumors, we analyzed Methylated DNA immunoprecipitation sequencing (MeDIP-Seq) data of a panel of LuCaP PDX tumors with high or low HOXB13 expression (**Fig.S5A**). Importantly, there were remarkably increased methylations near two CpG islands within the HOXB13 gene in the CRPC tumors with low HOXB13 expression (**Fig.5G**). In addition, analyses of human PCa samples demonstrated significantly negative correlations between HOXB13 and epigenetic silencers DNMT3A and EZH2, which are respectively DNA methyltransferases that catalyze 5-methylcytosine methylation and histone methyltranferases that recruit DNMT3A and facilitate DNA methylation^50^ (**Fig.S5B**). Moreover, overexpression of DNMT3A or EZH2 indeed inhibited HOXB13 expression in LNCaP cells (**Fig. 5H-I**).

### HOXB13 loss promotes PCa cell motility *in vitro* and xenograft tumor metastasis *in vivo*

Aberrant lipogenic programs have been shown to induce metastatic PCa^2, 3, 7^ and targeting of *de novo* lipogenesis exhibited therapeutic effects in preclinical models of CRPC^4, 51^. We thus hypothesized that HOXB13 plays a previously unknown role in regulating PCa metastasis through modulation of lipid accumulation. To test this hypothesis, we first examined HOXB13 regulation of PCa cell motility *in vitro*. As previous studies have reported a major role of HOXB13 in regulating cell growth, primarily through its acting as an AR cofactor^16, 19^, we also performed colony formation assay in parallel to control for cell motility caused by different growth rates. We found that androgen-dependent growth of LNCaP cells was abolished upon HOXB13 KD, which made it unsuitable for the study of cell motility (**Fig. S6A**). The growth of the CRPC cell line C4-2B is much less dependent on HOXB13 expression and the dependency was further reduced in cells pre-treated with AR antagonist Enzalutamide (**Fig. S6B**). Importantly, HOXB13 depletion significantly increased C4-2B cell invasion, which was rescued (i.e., repressed) by re-expression of WT, but not G84E, HOXB13 (**Fig. 6A**). Further analysis of the AR-negative PC-3 cells demonstrated that HOXB13 depletion remarkably increased cell invasion and migration, but did not show a major effect on PC-3 cell growth (**Fig. 6B & S6C-D**).

**Figure 6.**
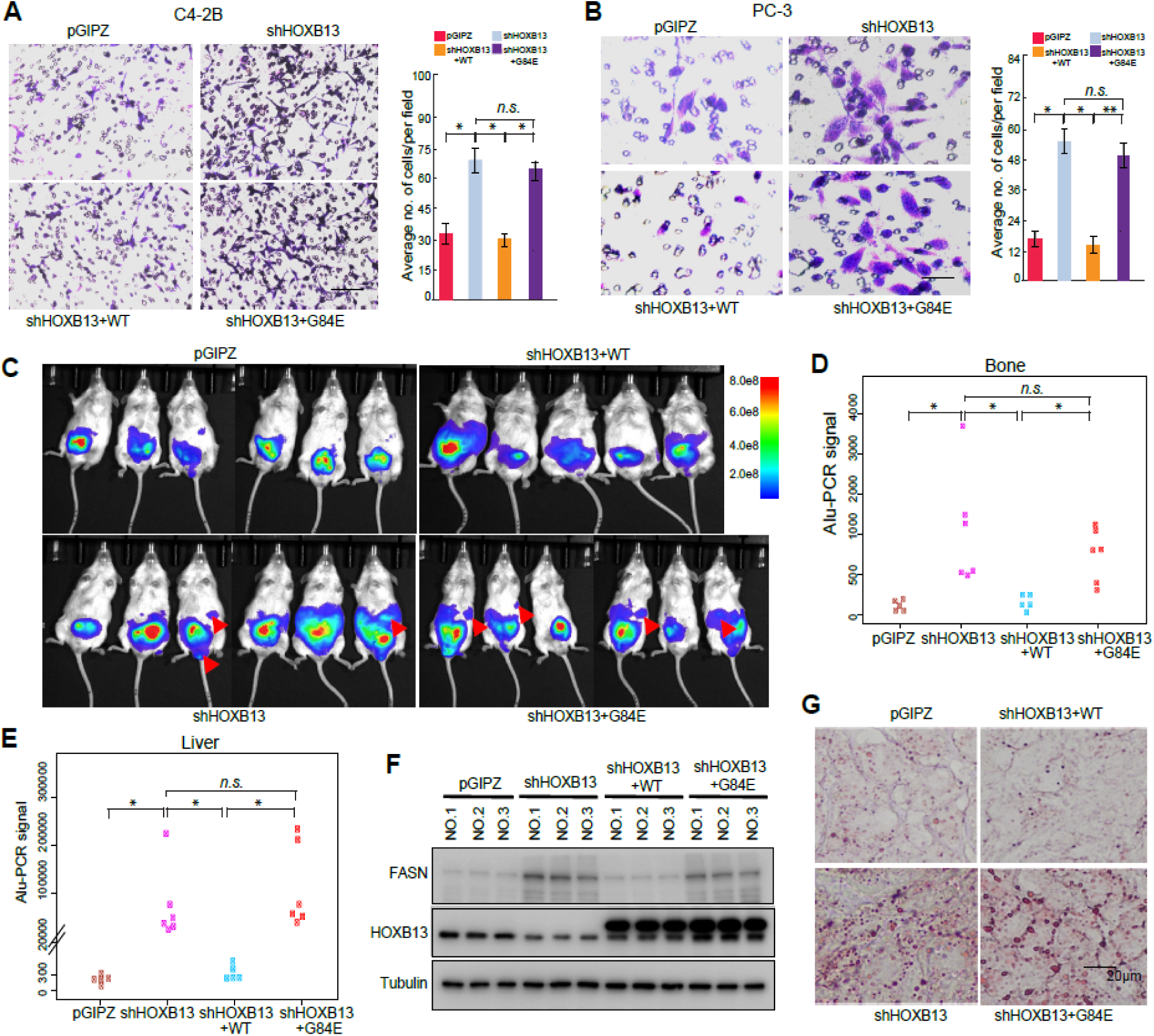
HOXB13 loss induces PCa cell motility *in vitro* and tumor metastasis *in vivo*. **A-B.** Cell invasion assays of C4-2B (**A**) and PC-3 (**B)** cells with shHOXB13 and/or rescue by HOXB13 WT or G84E mutant. Representative images are shown (left panels) and the number of invaded cells quantified (right panel). *\*\*p*<0.01, * *p*<0.05, *n.s.* not significant by Student’s *t*-test. **C**. IVIS live mouse imaging of orthotopic PC-3M xenograft tumors at three weeks after intra-prostate inoculation. Luciferase-labeled PC-3M cells with control (pGIPZ), shHOXB13, and shHOXB13 with HOXB13 WT or G84E re-expression were injected into the anterior prostates of nude SCID mice. Red arrows indicate local metastasis. Heatmap show the scale of IVIS signal intensity. **D-E**. Genomic DNA was isolated from mouse bone or liver and analyzed for metastasized PC-3M xenograft tumor cells by quantifying human Alu sequence by PCR. Y-axis shows human Alu signal detected by qPCR. \**p*<0.05, *n.s.* not significant, by Student’s *t*-test. **F.** PC-3M xenograft prostate tumors from three mice per group were collected and subjected for WB analysis. Tubulin is a loading control. **G.** Oil Red O staining of lipids in representative PC-3M xenograft prostate tumors.

To examine the effects of HOXB13 on PCa metastasis *in vivo*, we utilized the PC-3M cell line, which has high rates of metastasis after intraprostatic implantation^52^. Luciferase-labeled PC-3M cells with control, shHOXB13, or KD with WT or G84E re-expression were inoculated into the anterior prostates of nude SCID mice. Intra-prostatic xenograft tumors were observed in all mice after two weeks of inoculation with comparable initial tumor volume across groups (**Fig. S6E**). After three weeks of inoculation, regional metastases were observed in 2/6 mice with HOXB13-KD cells and 4/6 mice with HOXB13 KD and concurrent G84E re-expression, while mice in the control and WT-rescued groups did not show a clear sign of metastasis (**Fig. 6C**). As a control, the growth rates of primary tumors in the prostate were not significantly different across groups (**Fig. S6F**), in agreement with our *in vitro* data. Quantification of human Alu sequences by real-time PCR (Alu-qPCR) of liver and bone tissues at the endpoint revealed significantly more metastasis of xenograft tumors in the groups with HOXB13 KD or KD with G84E re-expression, as compared to the control group and the group with HOXB13 KD rescued by WT (**Fig. 6D-E**). *Ex Vivo* imaging of endpoint liver tissues for luciferase signals emanated from PC-3M cells confirmed that depletion of HOXB13 significantly promoted PC-3M tumor metastasis, which was abolished by the re-expression of WT, but not G84E, HOXB13 (**Fig. S6G**). WB analyses of representative xenograft prostate tumors confirmed increased FASN expression following HOXB13 KD *in vivo*, which was reduced by re-expressing WT, but not G84E, HOXB13 (**Fig. 6F**). Moreover, ORO staining showed greatly increased lipid accumulation in the HOXB13 KD group and the G84E re-expression group, being consistent with the higher metastatic rates of these tumors (**Fig. 6G)**. Taken together, these results indicate that HOXB13 loss promotes PCa cell migration and invasion *in vitro* and drives xenograft tumor metastasis *in vivo*.

### Therapeutic targeting of HOXB13-low tumors with FASN inhibitors

To investigate the clinical relevance of HOXB13 regulation of lipogenic pathways in human PCa, we examined FASN expression in publically available PCa expression profiling datasets and observed continuously increased expression from benign prostate to localized PCa to metastatic PCa (**Fig. S7A**). As FASN is a critical AR-induced gene^6^, its initial up-regulation in localized PCa may be driven by AR, while its further up-regulation in mCRPC may be attributed to HOXB13 loss. To confirm this at protein levels, we performed IHC of FASN using TMAs of human PCa tissues and found FASN protein mainly localized in the cytoplasm (**Fig. 7A**). In contrast to reduced HOXB13 staining in mCRPC relative to primary PCa, FASN expression was increased upon disease progression, being consistent with mRNA data (**Fig.7B**). Furthermore, analysis of FASN staining intensity in mCRPC with low HOXB13 (none or weak staining) and high HOXB13 (moderate or intense staining) revealed much stronger FASN staining in HOXB13-low tumors (**Fig. 7C**). About 25.5% HOXB13-low mCRPC tumors, compared to 15% of HOXB13-high tumors, showed strong FASN staining. These results confirm FASN up-regulation in mCRPC, especially those with low HOXB13, and support FASN as a clinically relevant therapeutic target.

**Figure 7.**
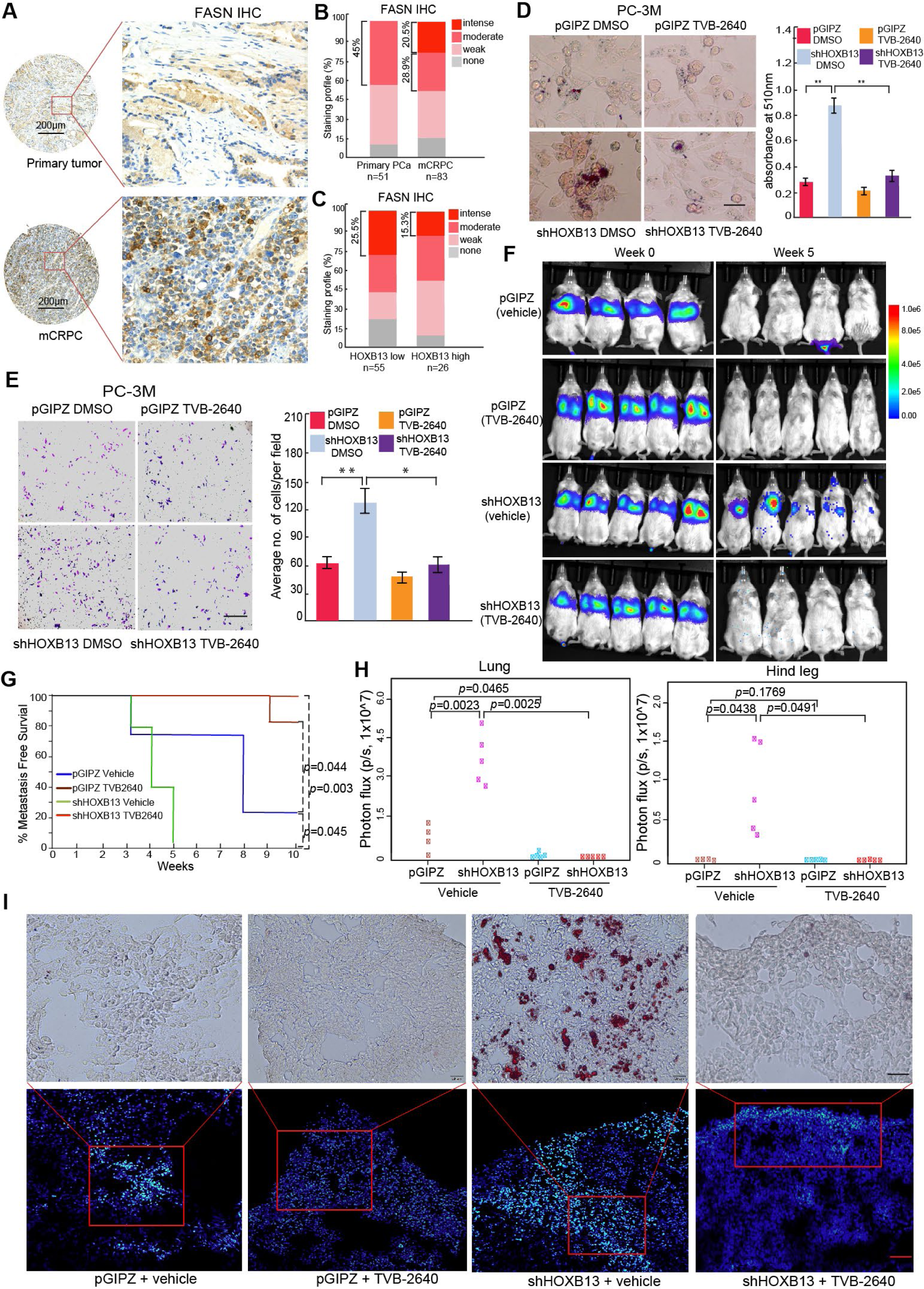
Therapeutic targeting of HOXB13-low tumors with FASN inhibitors. **A.** Immunohistochemistry (IHC) analyses of human primary PCa and mCRPC. Representative images of FASN staining in primary tumors (top panels) and metastatic CRPC (bottom panels) are shown in low-(×40) and high-magnification (×200). **B.** Quantification of FASN staining intensities in human PCa. Y axis shows percentage of tumors with none, weak, moderate, and intense IHC staining. **C.** Percentage of weak to strong FASN IHC staining in mCRPC tumors with low (none or weak) or high (moderate to intense) HOXB13 staining. IHC of FASN and HOXB13 were performed using adjacent tissue sections. **D.** Representative images of Oil Red O staining (left) and quantification (right) of lipid accumulation in HOXB13 KD PC-3M cells treated with DMSO or 5μM TVB-2640 for 3 days. Scale bar of 50 µm. Data shown are mean (±SEM) of triplicate wells. **p<0.01, *p<0.05, n.s. not significant by Student’s t-test. **E.** Cell invasion assays of control or HOXB13-KD PC-3M cells treated with DMSO or 5μM FASN inhibitor TVB-2640 for 3 days. Representative images are shown (left panels) and the number of invaded cells quantified (right panels). **p<0.01, *p<0.05, n.s. not significant by Student’s t-test. **F.** IVIS live mouse imaging of intravenous PC-3M xenograft tumors at week 0 and 5 after inoculation. Luciferase-labeled pGIPZ or shHOXB13 PC-3M cells were intravenously injected through tail vein of nude SCID mice. At 10 days after inoculation, mice were randomized to receive vehicle (30% PEG400) or TVB-2640 (100mg/Kg) every day. Heatmap shows IVIS signal intensity color scale. **G.** Kaplan-Meier analyses of metastasis-free survival of pGIPZ and HOXB13 KD mice treated with vehicle or TVB-2640. Metastasis was defined as whole-mouse IVIS signal higher than 1x10^5 Photons/s after 3 weeks of PCa cell inoculation. *P* values were determined by log rank test. **H.** *Ex Vivo* IVIS analysis and quantification of PC-3M tumor metastasis to the lung and hind leg. At the endpoint, lung and hind legs were collected and measured by *Ex vivo IVIS*. Y-axis shows the normalized luciferase intensity. Indicated *p*-values were by Student’s t-test. **I.** Oil Red O staining analysis of lipid accumulation in metastatic tumors. Tumors in the lung were identified by GFP (Green) using IF (bottom row) and adjacent section was used for Oil Red O staining (top). IF images shown here are 20x and Oil Red O staining images 40x magnification.

Next we tested whether pharmaceutical targeting of FASN blocks the oncogenic roles induced by HOXB13 loss in PCa cells using TVB-3166 or TVB-2640, orally available, potent, and selective FASN inhibitors that are also active in mice ^53^ and TVB-2640 is currently in phase II clinical trial for breast cancer and phase I trial for colorectal cancer. ORO staining demonstrated that TVB-3166 and TVB-2640 significantly reduced lipid accumulation in HOXB13-depleted PC-3M and C4-2B (**Fig. 7D & S7B**). In concordance with this, TVB-2640 significantly decreased HOXB13-KD PC-3M cell invasion and similar results were observed in C4-2B cells with HOXB13 depletion (**Fig.7E & S7C**). To evaluate the efficacy of FASN inhibitors in targeting HOXB13-mediated metastasis *in vivo*, we generated intravenous PCa xenograft models by inoculating luciferase-labeled PC-3M cells with control or HOXB13 KD to nude SCID mice through the tail vein (**Fig.7F, week 0**). At 10 days after PCa cell inoculation, mice with control or KD cells were each randomized to receive treatment with vehicle (30% PEG400) or FASN inhibitor TVB-2640 (100mg/kg, once daily) for 6 weeks by oral gavage. *In vivo* imaging system (IVIS) showed clear metastasis in HOXB13-KD mice at 5 weeks post inoculation, which was fully mitigated by TVB-2640 treatment (**Fig.7F, week 5**). In contrast, the control PC-3M cells did not show detectable whole mice IVIS signal. Further, TVB-2640 treatment did not apparently affect mouse body weight, indicating reasonable tolerability (**Fig. S7D**). Under vehicle-treated conditions, mice with HOXB13 KD cells showed significantly (*p*=0.045) reduced metastasis-free survival as compared to mice inoculated with pGIPZ control cells. Importantly, TVB-2640 treatment significantly prolonged metastasis-free survival of both pGIPZ mice (*p*=0.044) and HOXB13-KD mice (*p*=0.003) (**Fig. 7G**). *Ex vivo* IVIS analyses of dissected organs at the endpoint revealed significantly more HOXB13-KD PCa cell metastasis to the lung, hind leg, liver, and rib than the control cells, while TVB-2640 treatment abolished tumor metastasis *in vivo* (**Fig. 7H & S7E-G**). Accordingly, ORO staining showed massive lipid accumulation in metastasis tumor in lung of mice inoculated with HOXB13-KD cells, which was eliminated by TVB-2640 treatment (**Fig.7I**). In aggregates, our results demonstrated that FASN inhibitors may be useful to inhibit HOXB13-low PCa cell invasion and tumor metastasis by targeting lipid synthesis.

## DISCUSSION

HOXB13 is a critical transcription factor that is almost exclusively expressed in the prostate^14, 15^. Current understanding of HOXB13 is limited largely to its role in regulating androgen-dependent AR signaling and PCa growth^16, 18, 19, 24^. However, this role of HOXB13 failed to explain the well-noted association of germline G84E mutation of HOXB13 with early onset PCa^23^, as G84E did not increase HOXB13 interaction with AR, nor did it enhance AR signaling. Here we report a novel interaction between HOXB13 and HDAC3-NCoR co-repressor complex that suppresses the expression of PSA and many other lipid regulators. This interaction is disrupted by G84E mutation on HOXB13, leading to increased PSA levels. We speculate that PSA elevation through this AR-independent mechanism might add on to the PSA increase caused by aberrant AR signaling in cancerous cells, accounting for an earlier diagnosis of PCa patients with G84E mutation. Of noteworthy, previous studies have reported a lack of association of HOXB13 G84E mutation with Gleason score, tumor grade, and metastasis at diagnosis^54^, which is consistent with the fact that the mutation does not alter HOXB13 interaction with AR, the key driver of PCa growth. Our mechanistic study thus provides a molecular explanation of this dogma that HOXB13 G84E mutation is associated with early onset disease but NOT tumor aggressiveness. Although the difference was not significant, there was a better PCa-specific survival for HOXB13 G84E carriers^54^, which is in contrast to our findings of impaired ability of G84E to suppress lipid metabolism, consequently increasing metastasis of CRPC tumors. We reason that this may be due, at least in part, to earlier PCa diagnosis in G84E carriers, leading to earlier disease management. To clarify this, future analyses of HOXB13 G84E association with clinical outcomes in treatment- and age-controlled groups will be needed.

In the present study, we found that HOXB13 directly interacts with HDAC3 to recruit NCoR, which activates the histone deacetylase function of HDAC3 to remove histone acetylation at the regulatory elements of lipogenic genes. This complex is present in both AR+ LNCaP cells and AR-PC-3 cells, indicating its independence of AR. However, HOXB13 cistrome in the LNCaP cells would inevitably be affected by AR, FOXA1, and other luminal transcription factors^18^, either directly through protein-protein interaction or indirectly via alterations in chromatin accessibilities. This might account for the different HOXB13 binding sites between LNCaP and PC-3 cells that we have observed. Conversely, it has been previously reported that HOXB13 inhibits AR binding at some regulatory elements, indirectly resulting in suppression of gene expression^16^. For instance, the increased expression of PSA and FASN in HOXB13-KD LNCaP cells may be partially accounted by increased activities of AR due to HOXB13 depletion, as AR is a major promoter of lipogenesis^6^. Our study does not exclude the presence of such secondary mechanisms. However, by demonstrating the ability of HOXB13 to inhibit lipogenic programs in both LNCaP and PC-3 cells, we present a central model wherein HOXB13 directly suppresses lipid gene expression independently of AR, through actively recruiting HDAC3-NCoR for epigenetic reprogramming.

Although HOXB13 is known as a pioneer factor and a critical cofactor of AR and ARv7^17–19^, there has, surprisingly, been great controversy in the literature regarding its role in regulating PCa progression. There is an overall trend of HOXB13 in inducing AR-positive PCa cell growth^13, 16, 18, 19^, which however is not without exceptions^21, 22^. Understanding of HOXB13 function in AR-negative cells is even more limiting. Here, we clarify that the androgen-sensitive cell line LNCaP is strongly dependent on HOXB13 for growth, CRPC lines such as C4-2B is much less dependent, and AR-negative cell line PC-3 is independent of HOXB13 (**Fig. S6A-C**). These findings are consistent with the observed up-regulation of HOXB13 in PCa relative to benign tissue. HOXB13 becomes down-regulated as disease progresses from localized PCa to CRPC to NEPC, as the tumors gain metastatic features and become less dependent on AR signaling. At this stage, the function of HOXB13 to suppress lipid accumulation, cell invasion, and metastasis, as reported in this study, becomes apparent. Dissecting this new role of HOXB13 requires careful control of confounding effects caused by AR-dependent cell growth. Through orthotopic implantation of the AR-negative but HOXB13-positive PC-3M cells, here we clarify that HOXB13 depletion increases PC-3M xenograft tumor metastasis without clearly affecting the growth rate of its primary tumors. Such complex roles of HOXB13 in regulating PCa growth and metastasis should be carefully considered along with anti-androgen treatment histories when evaluating the association of HOXB13 expression with clinical outcomes of PCa patients.

Although there are increasing evidences associating aberrant lipid metabolism with PCa^2, 7^, the molecular mechanisms that drive lipogenic reprogramming has only begun to unveil. Here we report HOXB13 loss as a critical mechanism for hyperactive lipid biosynthesis and fatty acid accumulation in late-stage PCa. TMA analysis identified 30% CRPC tumors with low HOXB13 expression that may benefit from therapeutic targeting of its downstream pathways. Out of HOXB13-low CRPC, approximately 55% tumors showed moderate to strong FASN immunostaining, providing an avenue for targeting using FASN inhibitors. Multiple pharmaceutical inhibitors of FASN have been developed and tested in the recent years, including IPI-9119^4^, TVB-3166, and TVB-2640^55^. Preclinical studies of IPI-9119 showed a reduction in ARv7-driven CRPC growth and enhanced efficacy of enzalutamide in combinatorial treatment, but this compound is not available for clinical use^4^. The TVB series of compounds are under active preclinical and clinical development and presently TVB-2640 is in clinical trials in multiple cancer types including breast, lung, and colon cancers (clinicaltrials.gov). Here we demonstrate that TVB-2640 treatment abolishes HOXB13 loss-induced lipid accumulation, cell invasion, and xenograft tumor metastasis. It will be of great interest for future studies to further examine the efficacy of TVB-2640 in additional PCa preclinical models, either as a single agent or in combination with AR pathway inhibitors such as abiraterone, enzalutamide, or apalutamide, which will pave the way to a clinical trial of TVB-2640 for PCa, especially for those driven by HOXB13 loss.

## Materials and Methods

### Cell lines, chemical reagents, and antibodies

Prostate cancer cell lines LNCaP, C4-2B, 22Rv1, PC-3 and human embryonic kidney cell line HEK293T cells were obtained from American Type Culture Collection (ATCC) and cultured in either RPMI1640 or Dulbecco’s modified Eagle’s medium with 10% fetal bovine serum (FBS), 1% penicillin and streptomycin. Cells were authenticated, free of mycoplasma. Oil Red O (O0625) and TVB-3166 (SML1694) were purchased from Sigma, TVB-2640 (Catalog No. T15271) was from TargetMol. All antibodies used in this study are listed in **Supplementary Table 1.**

### Constructs and Lentivirus

HOXB13, HDAC1, HDAC3, EZH2 and NCOR1 constructs were first cloned into pCR8 Gateway compatible entry vector, and then transferred into pLenti-SFB, pLVX or pLenti6.3/V5 gateway compatible destination vector by LR clonase (Invitrogen). HOXB13 N-terminal and HDAC3 C-terminal truncates were generated by subcloning into modified pLV-EF1a-IRES-Puro vector (Biosettia). HOXB13 G84E, ΔMEIS, ΔHOX, WFQ-3A and NCOR1 ΔN1/2, ΔN1/2/3 and ΔDAD mutants were generated by Q5® Site-Directed Mutagenesis Kit (NEB, E0554S) with HOXB13 WT or NCOR1 WT as a template. The primers used in this study are listed in Supplementary Table 2 and all the plasmids were verified by sequencing. The pGIPZ lentiviral shRNA targeting 3′UTR of HOXB13 (Clone Id: V3LHS_403019) and control vector were purchased from Open Biosystems. SiRNA targeting AR was from Dharmacon (L-003400-00-0020). The shRNAs targeting HDAC1 (5′-CTATGGTCTCTACCGAAAA-3′) and HDAC3 (5′-GCATTGATGACCAGAGTTA-3′) were cloned into pLKO.1-TRC lentiviral vector (Addgene, #10878). For generation of lentivirus, HEK293T cells were transfected with psPAX2 and pMD2G at ratio 3:1. Supernatant containing lentiviruses was harvested at 48 h after transfection and filtered through a 0.45 μm filter. Lentiviruses, supplemented with 8 μg/mL polybrene, were used to infect PCa cells. Infected cells were selected with 2 μg/mL puromycin at 48 h after infection.

### Co-IP, WB, and chromatin fractionation assay

Co-IP experiments were carried out using standard protocol. Briefly, The whole cell lysates were extracted from trans transfected 293T cells or infected LNCaP cells or PC-3 cells by IP lysis buffer (50 mM Tris-Cl pH 7.4, 150 mM NaCl, 1 mM EDTA, 1% Triton X-100, Roche protease inhibitor cocktail). An aliquot of the cell lysate was kept as input for western blot analysis. Cell lysates were incubated with corresponding tag antibody at 4 °C overnight. For endogenous HOXB13 Co-IP in PC-3 cells, the extracted lysates were first pre-cleared with protein G-magnetic beads at 4 °C for 2 h followed by incubation with HOXB13 antibody (2 μg/sample, Santa Cruz, sc-28333) overnight. Dynabeads Protein A or G (Life Technologies), 25 µl per IP, were added the next day and incubated for 1 h at 4°C. The beads/protein complex was washed four times with IP lysis buffer and eluted with 30 µl 2× SDS sample buffer and subjected to western blot analysis using corresponding antibody.

For chromatin fractionation assay, chromatin was isolated as previously reported with the modifications^56^. Briefly, cells were resuspended in buffer A (10 mM HEPES, pH 7.9, 10 mM KCl, 1.5 mM MgCl2, 0.34 M sucrose, 10% glycerol, 1 mM DTT, 1× Roche protease inhibitor cocktail) and incubated on ice for 10 min. Then, final concentration of 0.1% Triton X-100 was added to cell suspension and vortexed for 15 s and spun down at 4 °C for 5 min at 1,000× g. The supernatant was kept as cytoplasmic fraction. Nuclei pellet was washed once with buffer A, and then resuspended in buffer B (3 mM EDTA, 75mM NaCl, 0.1% TritonX-100, 1 mM DTT, protease cocktail) for 30 min on ice. Nuclei was spun down for 5 min at 1,000× g at 4 °C and supernatant was saved as nuclear fraction. Insoluble chromatin was re-suspended in 1× SDS sample buffer.

### Mass spectrometry analysis

Mass spectrometry analysis were performed as described previously^57^. LNCaP cells stably expressing HOXB13 WT-SFB or G84E-SFB were lysed in NETN (100 mM NaCl, 20 mM Tris-Cl, pH 8.0, 1 mM EDTA, and 0.5% (vol/vol) NP-40) buffer containing protease inhibitors for 20 min at 4 °C. Crude lysates were subjected to centrifugation at 21,100 × g for 30 min. Supernatants were then incubated with streptavidin-conjugated beads (GE Healthcare) for 4 h at 4 °C. The beads were washed three times with NETN buffer, and bounded proteins were eluted with NETN buffer containing 2 mg/ml biotin (Sigma-Aldrich) for 1 h twice at 4 °C. The elutes were incubated with S-protein beads (EMD Millipore) overnight at 4 °C. The beads were eluted with SDS sample buffer and subjected to SDS-PAGE. Protein bands were excised and subjected to mass spectrometry analysis using Orbitrap Velos Pro™ system. The normalized spectral abundance factors were applied to calculate each detected protein to estimate relative protein levels^58^.

### RNA extraction, RT-qPCR, and RNA-Seq

RNA was extracted using the nucleospin RNA kit (Takara) according to the manufacturer’s recommended protocol. 500ng RNA was reverse transcribed into cDNA using the ReverTra Ace® qPCR RT Master Mix kit (Diagnocine) according to the manufacturer’s recommended protocol. QPCR was performed with 2xBullseye EvaGreen qPCR MasterMix (MIDSCI) and StepOne Plus (Applied Biosystems). All primers used here were listed in Supplementary Table 2. For RNA-seq, total RNA was isolated as described above. RNA-seq libraries were prepared from 0.5 μg high-quality DNA-free total RNA using NEBNext® Ultra RNA Library Prep Kit, according to the manufacturer’s instructions. The libraries passing quality control (equal size distribution between 250-400 bp, no adapter contamination peaks, no degradation peaks) were quantified using the Library Quantification Kit from Illiumina (Kapa Biosystems, KK4603). Libraries were pooled to a final concentration of 10nM and sequenced single-end using the Illumina HiSeq 4000.

### ChIP, ChIP-seq, and ATAC-seq

ChIP and ChIP-seq were performed using previously described protocol with following modifications^56^. LNCaP cells with control, HOXB13 KD, HOXB13 KD with WT or G84E rescue were cross-linked with 1% formaldehyde for 10 min at room temperature with gentle rotation and then quenched for 5 min with 0.125 M glycine. 5 million cells were used for each HOXB13 ChIP, 20-25 million cells were used for each HDAC1 or HDAC3 ChIP, 2 million cells were used for each H3K27ac ChIP. Chromatin was sonicated to an average length of 200-600 bp using E220 focused-ultrasonicator (Covaris). Supernatants containing chromatin fragments were pre-cleared with protein A agarose beads (Millipore) for 20 min and incubated with a specific antibody overnight at 4 °C on a nutator (antibody information were listed in Supplementary Table 1).then added 50 ul of protein A agarose beads and incubated for 2 h. Beads were washed twice with 1× dialysis buffer (2 mM EDTA, 50 mM Tris-Cl, pH 8.0) and four times with IP wash buffer (100 mM Tris-Cl, pH 9.0, 500 mM LiCl, 1% NP40, 1% Deoxycholate). The antibody/protein/DNA complexes were eluted with elution buffer (50 mM NaHCO3, 1% SDS), reversed the cross-links and DNA was purified with DNA Clean & Concentrator™-5 kit (ZYMO Research). For ChIP-qPCR, ChIPped DNA was diluted at 1:10 in water and 4µl diluted DNA was used as template for each qPCR reaction, ChIP-qPCR primers were used here were listed in Supplementary Table 2. ChIP-seq libraries were prepared from 3-5ng ChIPed DNA using NEBNext® Ultra™ II DNA Library Prep Kit (NEB, E7645S), according to the manufacturer’s instructions. Postamplification libraries were size selected at 250–450 bp in length using Agencourt AMPure XP beads from Beckman Coulter and were quantified using the Library Quantification Kit from Illiumina (Kapa Biosystems, KK4603). Libraries were pooled to a final concentration of 10nM and sequenced single-end using the Illumina HiSeq 4000.

ChIP for HOXB13, HDAC3 and H3K27ac in mCRPC LuCaP PDXs was performed as previously described^39^. Briefly, 50 mg of frozen tissue was cut into small pieces and homogenized using the BeadBug^TM^ benchtop homogenizer. The tissues were crosslinked in 2 steps with 2 mM of DSG (Pierce) for 10 minutes at room temperature followed by 1% Formaldehyde for 10 minutes. Crosslinked nuclei were then quenched with 0.125 M glycine for 5 minutes at room temperature and washed with PBS. Chromatin shearing, immunoprecipitation and libray preparation was performed as in prostate cancer cell line described above.

Transposase-Accessible Chromatin using sequencing (ATAC-seq) was performed as previously reported^59^. About 50,000 cells were collected, washed in PBS, pelleted by centrifugation and lysed and tagmented in 1x TD buffer, 2.5 μl Tn5 (Illumina), 0.01% Digitonin (Promega, G9441), 0.3x PBS in a 50 μl reaction volume. Samples were incubated at 37°C for 30 min at 300 rpm. Tagmented DNA was purified using the DNA Clean & Concentrator™-5 kit (ZYMO Research). Libraries were amplified as described previously^60^. Paired-end reads (50bp) were sequenced using an Illumina HiSeq 4000.

### Cell invasion and colony formation assay

Cell invasions assays were carried out as previously reported^52^. In brief, the cell suspension containing 300,000 (C4-2B) or 100,000 (PC-3) cells/mL in serum-free RPMI medium were prepared, 100 μL of cell suspension were transferred into the upper chamber. The lower chamber contained 500 μL of complete growth medium with 40% FBS. After incubation for 72 h, non-invading cells and matrigel were gently removed using a cotton-tipped swab. The inserts were fixed and stained for 15 min in 25% methanol containing 0.5% Crystal Violet. The images of invaded cells were captured under a bright-field microscope, and the number of invaded cells per field view was counted using the cell counter plugins in Image J. For colony formation assay, LNCaP (5x10^3^) and PC-3 (2x10^3^) cells per well were seeded on 12-well plate. After 2 weeks, the cells were first fixed by 4% paraformaldehyde and then stained with 0.05% crystal violet. The colonies were then imaged with ChemiDoc (BIO-RAD).

### Tissue acquisition and tissue microarray analysis

Tissue microarrays containing metastatic CRPC specimens were obtained as part of the University of Washington Medical Center Prostate Cancer Donor Program, which is approved by the University of Washington Institutional Review Board. All specimens for IHC were formalin fixed (decalcified in formic acid for bone specimens), paraffin embedded, and examined histologically for the presence of nonnecrotic tumor. TMAs were constructed with 1-mm diameter duplicate cores (*n* = 176) from CRPC patient tissues (*n* = 25 patients) consisting of visceral metastases and bone metastases (*n* = 88 sites) from patients within 8 hours of death. Two sets of TMAs of primary PCa (*n* = 51 patients, *n* = 51 sites) were used in this study. TMA (PCF 401334) was generated by the Northwestern University Pathology Core, and approved by the Northwestern University Institutional Review Board. TMA (CHTN_PrC_Prog1) was obtained through Cooperative Human Tissue Network (CHTN) at the University of Virginia.

Human TMA IHC staining was conducted using the Dako Autostainer Link 48 with enzyme-labeled biotin streptavidin system and the SIGMA*FAST* DAB Map Kit (MilliporeSigma). Antibodies used in IHC include anti-HOXB13 (1:400, 90944, Cell Signaling Technology) and anti-FASN (1:400, A301-323A, Bethyl). Images were captured with TissueFax Plus from TissueGnostics, exported to TissueFAX viewer, and analyzed using Photoshop CS4 (Adobe). HOXB13 and FASN immunostaining was scored blindly by a pathologist using a score of 0 to 3 for intensities of negative, weak, moderate or strong multiplied by percentage of stained cancer cells

### Murine orthotopic and intravenous xenograft studies

Mouse handling and experimental procedures were approved by the Center for Animal and Comparative Medicine in Northwestern University School of Medicine in accordance with the US National Institutes of Health Guidelines for the Care and Use of Laboratory Animals and the Animal Welfare Act. All LuCaP PDXs were derived from resected metastatic prostate cancer with informed consent of patient donors as described by previously^46^ under a protocol approved by the University of Washington Human Subjects Division IRB. NOD SCID male mice at 6-7 weeks old were purchased from Charles River Laboratories. Orthotopic implantations were carried out as previously described^52^. Briefly, a suspension of luciferase labeled PC-3M with control, HOXB13 KD, HOXB13 KD with WT or G84E re-expression stable cells (5× 10^5^ cells in 30 μl of PBS) was injected into the mouse anterior prostate. To monitor tumor growth and metastasis *in vivo*, D-Luciferin (100 μl of 15 mg/mL stock; Goldbio) was injected intraperitoneally into mice, bioluminescence was measured using Lago Bioluminescence/Fluorescence Imaging System at Northwestern core facility. For *Ex vivo* IVIS assay, at the endpoint, mice were euthanized 10min after injected D-Luciferin, then liver, lung and bone were collected and imaged immediately by Lago Bioluminescence/Fluorescence Imaging System. Alu-PCR analysis of tumor metastasis was performed as previously reported^61^.

For intravenous Xenograft model. NOD SCID male mice at 6-7 weeks old were purchased from Charles River Laboratories. A suspension of luciferase labeled PC-3M with control, HOXB13 KD, stable cells (2× 10^6^ cells in 200 μl of 1xPBS) were injected into mice through the tail vein. 10 days after inoculation, Mice were randomly divided into two groups and treated with vehicle (30% PEG400) or FASN inhibitor TVB-2640 (100mg/Kg) once daily for 6 weeks by oral gavage. Tumor metastasis was measured as described above by Lago Bioluminescence System. At the endpoint, liver, lung, hind leg and rib of mouse were collected and *Ex vivo* IVIS assay was performed as described above.

### Lipidomics analysis and oil red O staining of neutral lipids

Untargeted lipidomics analysis by Q-TOF LC/MS was done at Mass Spectrometry Core of University of Illinois at Chicago. Lipid molecules were identified and quantified with LipidSearch 4.1.9 software. The lipidomic analysis was performed as previously described (Chen et al., 2018). Briefly, 10 million LNCaP cells were harvested and washed three times with 1ml phosphate buffered saline (PBS) to remove any traces of culture medium. Nonpolar lipids were extracted using Folch method. The organic phase containing the nonpolar lipids was dried in a SpeedVac rotary evaporator with no heat. Lipid samples were reconstituted in 35 μl of 50% isopropanol (IPA)/50% MeOH. 10 μl of samples was subjected to Q-TOF LC/MS analysis. Oil Red O staining of prostate cancer cells, frozen LuCaP PDXs, frozen prostate Xenograft tumor and quantification of staining were performed using Lipid (Oil Red O) Staining Kit (Biovision, catalog # K580-24). For lipid staining of metastasis tumor in lung, tumor was identified by GFP and adjacent section was used for Oil Red O staining.

### Statistical Analysis

For each independent *in vitro* experiment, at least three technical replicates were used. Most *in vitro* experiments were repeated independently for three times and some were repeated twice. Two-tailed unpaired Student’s t tests were used to assess statistical significances in quantitative RT-PCR experiments and cell-based functional assays. Boxplot elements are defined as center line, median; box limits, upper and lower quartiles; whiskers, 1.5x interquartile range; points, outliers. One-way ANOVA was used to determine statistically significant differences across treatment groups in the xenograft studies. P less than 0.05 was considered statistically significant. Kaplan-Meier analyses of metastasis-free survival of mice were performed using log-rank test.

### RNA-seq and ChIP-seq analysis

RNA-seq reads were mapped to NCBI human genome GRCh38 using STAR version 1.5.2. Raw counts of genes were calculated by STAR. FPKM values (Fragments Per Kilobase of transcript per Million mapped reads) were calculated by in house perl script. Differential gene expression was analyzed by R Bioconductor DESeq2 package, which uses shrinkage estimation for dispersions and fold changes to improve stability and interpretability of estimates. ChIP-seq reads were aligned to the Human Reference Genome (assembly hg19) using Bowtie2 2.0.5. ChIP-seq peak identification, overlapping, subtraction and feature annotation of enriched regions were performed using HOMER (Hypergeometric Optimization of Motif EnRichment) suite.

Weighted Venn diagrams were created by R package Vennerable. Heatmap views of ChIP-seq were generated by deepTools. Raw data were uploaded to GEO as GSE153586.

#### Analysis of HOXB13 gene methylation in clinical PCa samples

HOXB13 methylation in primary PCa and benign prostate tissues in TCGA-PRAD methylation dataset was analyzed using the SMART App^48^. Alignment of methylation data relative to HOXB13 gene structure and nearby CpG islands was performed using the Wanderer tool^49^. HOXB13 gene methylation in CRPC tumors of LuCaP PDX models was determined by MeDIP-seq. Briefly, genomic DNA from the LuCaP PDXs was sheared using a Covaris Sonicator E220 and size selection performed with AMPure XP beads (Beckman Coulter) to retain 150-250 bp DNA fragments. MeDIP-seq was done following previously published methods^62^.

### Data and code availability

All sequencing data (RNA-seq and ChIP-seq) generated for the study has been deposited in GEO (GSE153586). This study did not systematically generate new code, software or algorithm.

All data analyses were performed using published software with the parameters indicated in the Methods section.

https://www.ncbi.nlm.nih.gov/geo/query/acc.cgi?acc=GSE153586

### Author Contributions

J.Y. and X.L. conceived the project and designed the experiments. J.C.Z., S.C.B., M.L.F., and J.Y. conducted bioinformatics and statistics analysis. F.W. and G.G. assisted with *in vivo* mouse experiment. K.F. performed IP-MS experiment and tissue microarray acquisition. J.R., X.L. and X.Y. performed HOXB13 and FASN IHC scoring. J.E.B. carried out MeDIP-seq experiment. E.C. provided LuCaP PDX tissues. X.L., J.C.Z., and J.Y. wrote the original manuscript. N.C., W.J.C., and M.L.F. consulted on the project and edited the manuscript.

## ACKNOWLEDGEMENTS

This work was partially supported by the Northwestern University Pathology Core Facility, Center for Advanced Microscopy/Nikon Imaging Center, and Robert H. Lurie Comprehensive Cancer Center Support Grant (NCI P30CA060553). NGS was done at the University of Chicago Genomic Facility and lipid profiling at Mass Spectrometry Core, University of Illinois at Chicago. IP-MS was done at Taplin Mass Spectrometry Facility of Harvard Medical School. Funding supports for the work include the NIH/NCI training grant T32CA09560 (to GG), prostate cancer SPORE P50CA180995 (to JY), R50CA211271 (to JCZ), R01CA257446 (to JY), and Prostate Cancer Foundation 2017CHAL2008 (to JY, JCZ). Generation and maintenance of the LuCaP PDX models were partially funded by NIH awards P50CA97186 and PO1CA163227.

## REFERENCE

1. Labbe, D.P. et al. High-fat diet fuels prostate cancer progression by rewiring the metabolome and amplifying the MYC program. Nat Commun 10, 4358 (2019).

2. Chen, J. et al. Compartmentalized activities of the pyruvate dehydrogenase complex sustain lipogenesis in prostate cancer. Nat Genet 50, 219–228 (2018).

3. Chen, M. et al. An aberrant SREBP-dependent lipogenic program promotes metastatic prostate cancer. Nat Genet 50, 206–218 (2018).

4. Zadra, G. et al. Inhibition of de novo lipogenesis targets androgen receptor signaling in castration-resistant prostate cancer. Proc Natl Acad Sci U S A 116, 631–640 (2019).

5. Poulose, N. et al. Genetics of lipid metabolism in prostate cancer. Nat Genet 50, 169–171 (2018).

6. Butler, L.M., Centenera, M.M. & Swinnen, J.V. Androgen control of lipid metabolism in prostate cancer: novel insights and future applications. Endocr Relat Cancer 23, R219–27 (2016).

7. Yue, S. et al. Cholesteryl ester accumulation induced by PTEN loss and PI3K/AKT activation underlies human prostate cancer aggressiveness. Cell Metab 19, 393–406 (2014).

8. Mitra, R., Chao, O., Urasaki, Y., Goodman, O.B. & Le, T.T. Detection of lipid-rich prostate circulating tumour cells with coherent anti-Stokes Raman scattering microscopy. BMC Cancer 12, 540 (2012).

9. Swinnen, J.V., Esquenet, M., Goossens, K., Heyns, W. & Verhoeven, G. Androgens stimulate fatty acid synthase in the human prostate cancer cell line LNCaP. Cancer Res 57, 1086–90 (1997).

10. Swinnen, J.V., Ulrix, W., Heyns, W. & Verhoeven, G. Coordinate regulation of lipogenic gene expression by androgens: evidence for a cascade mechanism involving sterol regulatory element binding proteins. Proc Natl Acad Sci U S A 94, 12975–80 (1997).

11. Han, W. et al. Reactivation of androgen receptor-regulated lipid biosynthesis drives the progression of castration-resistant prostate cancer. Oncogene 37, 710–721 (2018).

12. Berger, M.F. et al. Variation in homeodomain DNA binding revealed by high-resolution analysis of sequence preferences. Cell 133, 1266–76 (2008).

13. Huang, Q. et al. A prostate cancer susceptibility allele at 6q22 increases RFX6 expression by modulating HOXB13 chromatin binding. Nat Genet 46, 126–35 (2014).

14. Sreenath, T., Orosz, A., Fujita, K. & Bieberich, C.J. Androgen-independent expression of hoxb-13 in the mouse prostate. Prostate 41, 203–7 (1999).

15. Edwards, S. et al. Expression analysis onto microarrays of randomly selected cDNA clones highlights HOXB13 as a marker of human prostate cancer. Br J Cancer 92, 376–81 (2005).

16. Norris, J.D. et al. The homeodomain protein HOXB13 regulates the cellular response to androgens. Mol Cell 36, 405–16 (2009).

17. Economides, K.D. & Capecchi, M.R. Hoxb13 is required for normal differentiation and secretory function of the ventral prostate. Development 130, 2061–9 (2003).

18. Pomerantz, M.M. et al. The androgen receptor cistrome is extensively reprogrammed in human prostate tumorigenesis. Nat Genet 47, 1346–51 (2015).

19. Chen, Z. et al. Diverse AR-V7 cistromes in castration-resistant prostate cancer are governed by HoxB13. Proc Natl Acad Sci U S A 115, 6810–6815 (2018).

20. Augello, M.A. et al. CHD1 Loss Alters AR Binding at Lineage-Specific Enhancers and Modulates Distinct Transcriptional Programs to Drive Prostate Tumorigenesis. Cancer Cell 35, 603–617 e8 (2019).

21. Jung, C., Kim, R.S., Zhang, H.J., Lee, S.J. & Jeng, M.H. HOXB13 induces growth suppression of prostate cancer cells as a repressor of hormone-activated androgen receptor signaling. Cancer Res 64, 9185–92 (2004).

22. VanOpstall, C. et al. MEIS-mediated suppression of human prostate cancer growth and metastasis through HOXB13-dependent regulation of proteoglycans. Elife 9(2020).

23. Ewing, C.M. et al. Germline mutations in HOXB13 and prostate-cancer risk. N Engl J Med 366, 141–9 (2012).

24. Whitington, T. et al. Gene regulatory mechanisms underpinning prostate cancer susceptibility. Nat Genet 48, 387–97 (2016).

25. Mazrooei, P. et al. Cistrome Partitioning Reveals Convergence of Somatic Mutations and Risk Variants on Master Transcription Regulators in Primary Prostate Tumors. Cancer Cell 36, 674–689 e6 (2019).

26. Spisak, S. et al. CAUSEL: an epigenome- and genome-editing pipeline for establishing function of noncoding GWAS variants. Nat Med 21, 1357–63 (2015).

27. Heinzel, T. et al. A complex containing N-CoR, mSin3 and histone deacetylase mediates transcriptional repression. Nature 387, 43–8 (1997).

28. Abbas, A. & Gupta, S. The role of histone deacetylases in prostate cancer. Epigenetics 3, 300–9 (2008).

29. Horlein, A.J. et al. Ligand-independent repression by the thyroid hormone receptor mediated by a nuclear receptor co-repressor. Nature 377, 397–404 (1995).

30. Chen, J.D. & Evans, R.M. A transcriptional co-repressor that interacts with nuclear hormone receptors. Nature 377, 454–7 (1995).

31. Guenther, M.G., Barak, O. & Lazar, M.A. The SMRT and N-CoR corepressors are activating cofactors for histone deacetylase 3. Mol Cell Biol 21, 6091–101 (2001).

32. You, S.H. et al. Nuclear receptor co-repressors are required for the histone-deacetylase activity of HDAC3 in vivo. Nat Struct Mol Biol 20, 182–7 (2013).

33. Emmett, M.J. & Lazar, M.A. Integrative regulation of physiology by histone deacetylase 3. Nat Rev Mol Cell Biol 20, 102–115 (2019).

34. Knutson, S.K. et al. Liver-specific deletion of histone deacetylase 3 disrupts metabolic transcriptional networks. EMBO J 27, 1017–28 (2008).

35. Sun, Z. et al. Hepatic Hdac3 promotes gluconeogenesis by repressing lipid synthesis and sequestration. Nat Med 18, 934–42 (2012).

36. Feng, D. et al. A circadian rhythm orchestrated by histone deacetylase 3 controls hepatic lipid metabolism. Science 331, 1315–9 (2011).

37. Kent, L.N. & Leone, G. The broken cycle: E2F dysfunction in cancer. Nat Rev Cancer 19, 326–338 (2019).

38. Wang, S. et al. Target analysis by integration of transcriptome and ChIP-seq data with BETA. Nat Protoc 8, 2502–15 (2013).

39. Pomerantz, M.M. et al. Prostate cancer reactivates developmental epigenomic programs during metastatic progression. Nat Genet 52, 790–799 (2020).

40. Shang, Y., Myers, M. & Brown, M. Formation of the androgen receptor transcription complex. Mol Cell 9, 601–10 (2002).

41. Hodgson, M.C. et al. The androgen receptor recruits nuclear receptor CoRepressor (N-CoR) in the presence of mifepristone via its N and C termini revealing a novel molecular mechanism for androgen receptor antagonists. J Biol Chem 280, 6511–9 (2005).

42. Sun, Z. et al. Deacetylase-independent function of HDAC3 in transcription and metabolism requires nuclear receptor corepressor. Mol Cell 52, 769–82 (2013).

43. Guo, C. et al. Regulated clearance of histone deacetylase 3 protects independent formation of nuclear receptor corepressor complexes. J Biol Chem 287, 12111–20 (2012).

44. Chng, K.R. et al. A transcriptional repressor co-regulatory network governing androgen response in prostate cancers. EMBO J 31, 2810–23 (2012).

45. Wang, Z. et al. Genome-wide mapping of HATs and HDACs reveals distinct functions in active and inactive genes. Cell 138, 1019–31 (2009).

46. Nguyen, H.M. et al. LuCaP Prostate Cancer Patient-Derived Xenografts Reflect the Molecular Heterogeneity of Advanced Disease an--d Serve as Models for Evaluating Cancer Therapeutics. Prostate 77, 654–671 (2017).

47. Shroff, E.H. et al. MYC oncogene overexpression drives renal cell carcinoma in a mouse model through glutamine metabolism. Proc Natl Acad Sci U S A 112, 6539–44 (2015).

48. Li, Y., Ge, D. & Lu, C. The SMART App: an interactive web application for comprehensive DNA methylation analysis and visualization. Epigenetics Chromatin 12, 71 (2019).

49. Diez-Villanueva, A., Mallona, I. & Peinado, M.A. Wanderer, an interactive viewer to explore DNA methylation and gene expression data in human cancer. Epigenetics Chromatin 8, 22 (2015).

50. Vire, E. et al. The Polycomb group protein EZH2 directly controls DNA methylation. Nature 439, 871–4 (2006).

51. Watt, M.J. et al. Suppressing fatty acid uptake has therapeutic effects in preclinical models of prostate cancer. Sci Transl Med 11(2019).

52. Jin, H.J., Zhao, J.C., Ogden, I., Bergan, R.C. & Yu, J. Androgen receptor-independent function of FoxA1 in prostate cancer metastasis. Cancer Res 73, 3725–36 (2013).

53. Heuer, T.S. et al. FASN Inhibition and Taxane Treatment Combine to Enhance Anti-tumor Efficacy in Diverse Xenograft Tumor Models through Disruption of Tubulin Palmitoylation and Microtubule Organization and FASN Inhibition-Mediated Effects on Oncogenic Signaling and Gene Expression. EBioMedicine 16, 51–62 (2017).

54. Kote-Jarai, Z. et al. Prevalence of the HOXB13 G84E germline mutation in British men and correlation with prostate cancer risk, tumour characteristics and clinical outcomes. Ann Oncol 26, 756–761 (2015).

55. Ventura, R. et al. Inhibition of de novo Palmitate Synthesis by Fatty Acid Synthase Induces Apoptosis in Tumor Cells by Remodeling Cell Membranes, Inhibiting Signaling Pathways, and Reprogramming Gene Expression. EBioMedicine 2, 808–24 (2015).

56. Xu, B. et al. Altered chromatin recruitment by FOXA1 mutations promotes androgen independence and prostate cancer progression. Cell Res 29, 773–775 (2019).

57. Fong, K.W., Zhao, J.C., Song, B., Zheng, B. & Yu, J. TRIM28 protects TRIM24 from SPOP-mediated degradation and promotes prostate cancer progression. Nat Commun 9, 5007 (2018).

58. Paoletti, A.C. et al. Quantitative proteomic analysis of distinct mammalian Mediator complexes using normalized spectral abundance factors. Proc Natl Acad Sci U S A 103, 18928–33 (2006).

59. Corces, M.R. et al. An improved ATAC-seq protocol reduces background and enables interrogation of frozen tissues. Nat Methods 14, 959–962 (2017).

60. Buenrostro, J.D., Giresi, P.G., Zaba, L.C., Chang, H.Y. & Greenleaf, W.J. Transposition of native chromatin for fast and sensitive epigenomic profiling of open chromatin, DNA-binding proteins and nucleosome position. Nat Methods 10, 1213–8 (2013).

61. Song, B. et al. Targeting FOXA1-mediated repression of TGF-beta signaling suppresses castration-resistant prostate cancer progression. J Clin Invest 129, 569–582 (2019).

62. Shen, S.Y., Burgener, J.M., Bratman, S.V. & De Carvalho, D.D. Preparation of cfMeDIP-seq libraries for methylome profiling of plasma cell-free DNA. Nat Protoc 14, 2749–2780 (2019).

