## Supplemental Figures for "HOXB13 suppresses *de novo* lipogenesis through HDAC3-mediated epigenetic reprogramming"

**Contents**:

**Figure S1**. Genome-wide analysis revealed an essential molecular function of HOXB13 in suppressing *de novo* lipogenesis.

**Figure S2**. HOXB13 protein interacts with HDAC3 protein.

**Figure S3**. HOXB13 recruits HDAC3 to target chromatin to catalyze histone deacetylation.

**Figure S4**. HDAC3 is required for HOXB13-mediated suppression of de novo lipogenesis.

**Figure S5**. HOXB13 is hypermethylated and down-regulated in CRPC.

**Figure S6**. HOXB13 loss promotes PCa cell motility *in vitro* and xenograft tumor metastasis *in vivo.*

**Figure S7**. Therapeutic targeting of HOXB13-low tumors with FASN inhibitors.

**Supplementary Table 1**. Antibodies that were utilized in this study.

**Supplementary Table 2.** Oligonucleotides that were used in this study.

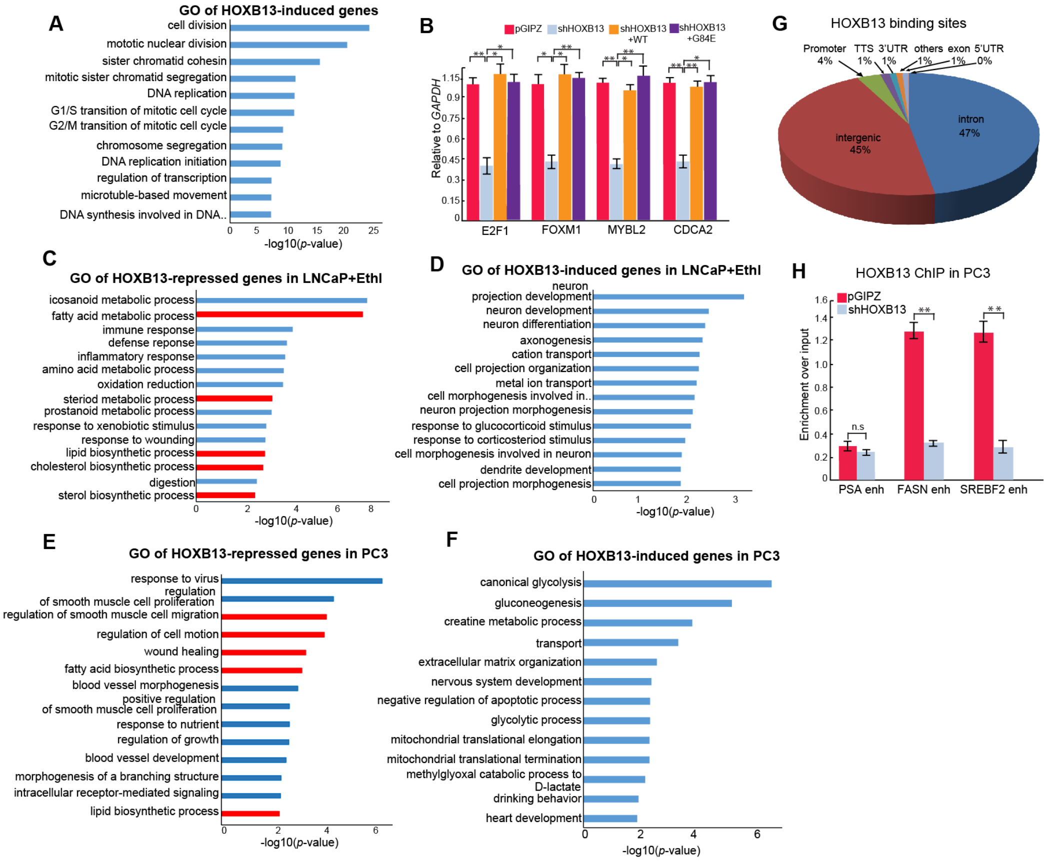


**Figure S1. Genome-wide analysis revealed an essential molecular function of HOXB13 in suppressing *de novo* lipogenesis.**

**A**. GO analysis of HOXB13-induced genes identified in Fig. 1A. Top enriched molecular concepts are shown on Y-axis, while X-axis indicates enrichment significance.

**B**. RT-qPCR validation of key cell cycle gene regulation by HOXB13 in LNCaP cells. Data were normalized to GAPDH. Data shown are mean (±SEM) of technical replicates from one representative experiment of three. Two-tailed unpaired Student's t-test, *$p < 0.05$, **$p < 0.01$, *ns*, not significant.

**C-D**. GO analysis of HOXB13-repressed (**C**) and -induced genes (**D**) in hormone-deprived LNCaP cells. LNCaP cells were infected with control shRNA (pGIPZ) or shRNA targeting HOXB13 (shHOXB13) for 4 days followed by hormone starvation for 3 days. Cells were harvested for RNA-seq analysis in replicate experiments. HOXB13-induced and -repressed genes were derived by comparing control cells with HOXB13-KD cells using FDR<0.05 and fold change >2.0. GO analysis was performed using DAVID, Top enriched molecular concepts are shown. X-axis indicates enrichment significance.

**E-F**. GO analysis of HOXB13-repressed (**E**) and -induced genes (**F**) in AR-negative PC-3 cells. PC-3 cells were subjected to control or shHOXB13 for 7 days and then harvested for RNA-seq in replicate experiments. HOXB13-repressed genes were derived by comparing control cells with HOXB13-KD cells using FDR<0.05 and fold change >=2. GO analysis was performed using DAVID, Top enriched molecular concepts are shown. X-axis indicates enrichment significance.

**G**. Pie Chart showing the genomic distribution of HOXB13 binding sites in LNCaP cells.

**H**. ChIP-qPCR analyses of H3K27ac at lipogenic gene enhancers in PC-3 cells with control or HOXB13 KD. *$p < 0.05$, **$p < 0.01$, *ns*, not significant.

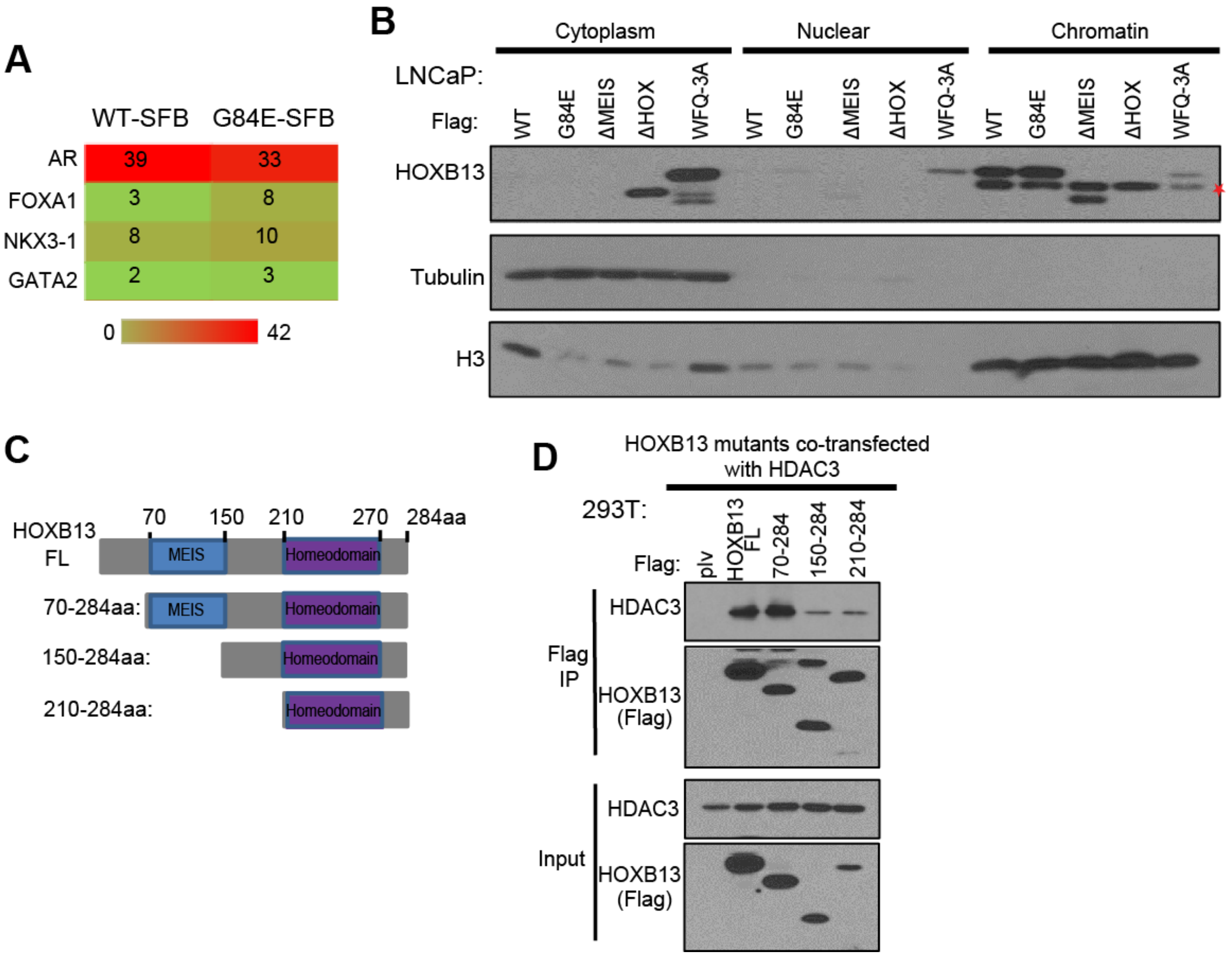


**Figure S2. HOXB13 protein interacts with HDAC3 protein.**

**A**. Heatmap showing AR co-factors that were enriched by HOXB13-WT or G84E mutant in a Mass Spec experiment.

**B**. Fractionation assay showing cellular localizations of HOXB13 WT and mutants in LNCaP cells. Asterisk indicates endogenous HOXB13.

**C-D**. Schematic illustration of a series of HOXB13 deletion mutants (**C**) and their interaction with HDAC3 (**D**). Whole cell lysates from 293T cells co-transfected with HA-HDAC3 along with Flag-tagged HOXB13 full-length (WT) or its deletion mutants were subjected to co-IP using an anti-Flag antibody.

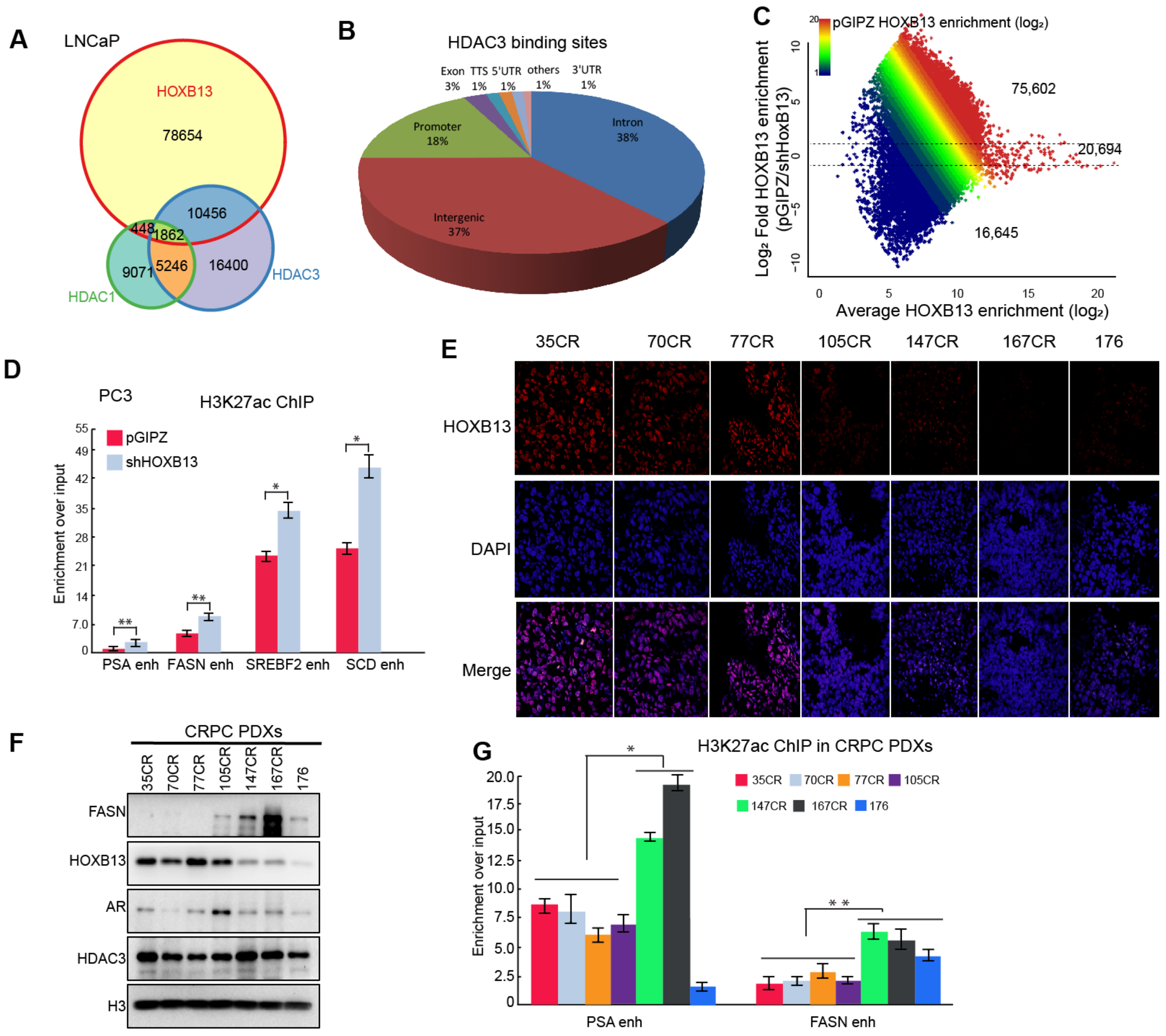


**Figure S3. HOXB13 recruits HDAC3 to target chromatin to catalyze histone deacetylation.**
**A**. Venn diagram showing overlap of HOXB13, HDAC1 and HDAC3 binding sites in LNCaP.
**B**. Pie Chart showing the genomic distribution of HDAC3 binding sites in LNCaP cells.
**C**. MA plot showing differential HOXB13 ChIP-seq enrichment in control (pGIPZ) and HOXB13 KD (shHOXB13) LNCaP cells. Color encodes the intensity of HOXB13 ChIP-seq in control cells and dotted lines for 2-fold differences.
**D**. ChIP-qPCR analyses of H3K27ac at lipogenic gene enhancers in PC3 cells with control or HOXB13 KD. *$p < 0.05$, **$p < 0.01$, *ns*, not significant.
**E**. Immunofluorescence showing HOXB13 staining in 7 mCRPC PDXs. Representative images of HOXB13 staining are shown in high-magnification (×60). DAPI staining marks nuclei.
**F**. WB analysis of the expression of HOXB13, FASN, AR, and HDAC3 in mCRPC PDXs.
**G**. ChIP-qPCR analysis of H3K27ac at PSA and FASN enhancer in mCRPC PDXs. Student's t-test, *$p < 0.05$, **$p < 0.01$, *ns*, not significant.

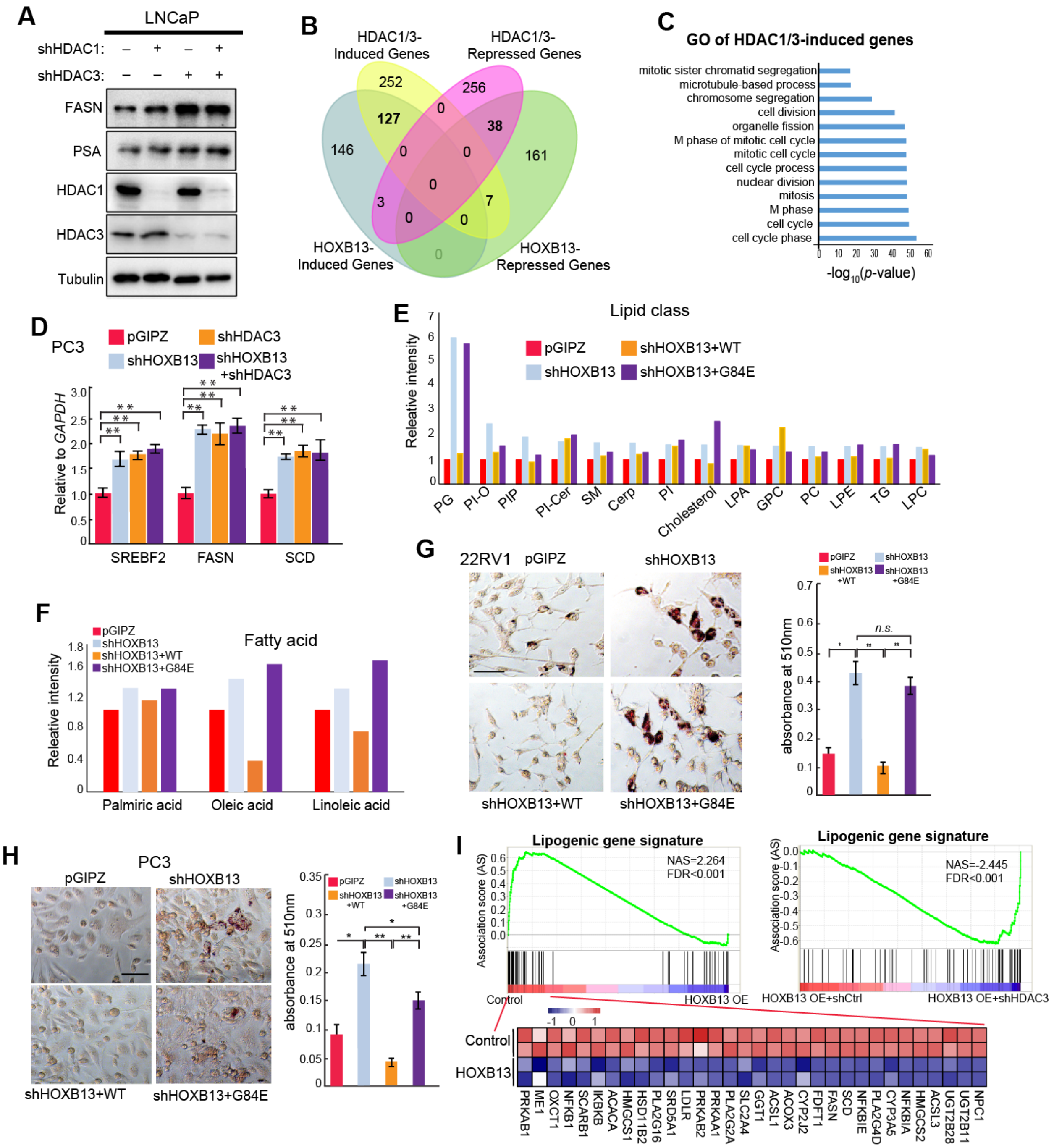

**Figure S4. HDAC3 is required for HOXB13-mediated suppression of *de novo* lipogenesis.**
**A**.WB analysis of FASN and PSA in LNCaP cells with KD of HDAC1, HDAC3 or both. Whole cell lysates from cells with indicated knockdown were analyzed by WB using anti-FASN, anti-PSA, anti-HDAC1 and anti-HDAC3 antibody, Tubulin was used as a loading control.
**B**. Venn Diagram showing overlap between HOXB13- or HDAC1/3-induced or –repressed genes in LNCaP cells.
**C**. GO analysis of HDAC1/3-induced genes found in LNCaP cells showing the top enriched molecular concepts.

**D**. RT-qPCR analysis of SREBF2, FASN, and SCD mRNA levels in PC-3 cells with KD of HOXB13, HDAC3, or both. RT-qPCR data were normalized to GAPDH. Data shown are mean (±SEM) of technical replicates from one representative experiment of three. Two-tailed unpaired Student's t-test, $^{*}p < 0.05$, $^{**}p < 0.01$.
**E**. Untargeted lipidomics analysis of LNCaP cells with control, HOXB13 KD, KD with concomitant rescue by HOXB13 WT or G84E. Lipids were extracted using Folch method and the extracted lipids were subjected to untargeted lipidomics analysis by Q-TOF LC/MS. Top 14 HOXB13-repressed lipid class are shown. PG, Glycerophosphoglycerols; PI-O, 1-alkyl,2-acylglycerophosphoinositols; PIP, Glycerophosphoinositol monophosphates; PI-Cer, Ceramide phosphoinositols; SM, Sphingomyelins; Cerp, Ceramide 1-phosphates; PI, Glycerophosphoinositols; Cholesterol; LPA, lysoglycerophosphates; GPC, Glycanglycerophosphocholines; PC, Glycerophosphocholines; LPE, lysophosphatidylethanolamine; TG, Triradylglycerolipids; LPC, lysoglycerophosphocholines.
**F**. Analysis of fatty acids in LNCaP cells with control, HOXB13 KD, and KD with concomitant re-expression of WT or G84E HOXB13.
**G-H**. Oil Red O staining and quantification of lipid accumulation in 22Rv1 (**G**) and PC-3 (**H**) with shHOXB13 and rescue by WT or G84E HOXB13. Data (left) show are representative images of Oil Red O staining with *Scale bar* of 50 µm. The stain was extracted in isopropanol and quantified at 510 nm (right). Data shown are mean (±SEM) of triplicate wells. $^{**}p<0.01$, $^{*}p<0.05$, *n.s.* not significant, by two-tailed unpaired Student's *t*-test.
**I**. GSEA of lipogenic gene signature in LNCaP cells with control (pLKO+plv) or HOXB13 OE (**left**) or HOXB13-overexpressing cells with control (shCtrl) or HDAC3 KD (**right**). The lipogenic gene signature was compiled from the lipid metabolism pathways shown in bold in Fig. 4J. Shown at the bottom are heatmaps of GSEA leading-edge gene expression.

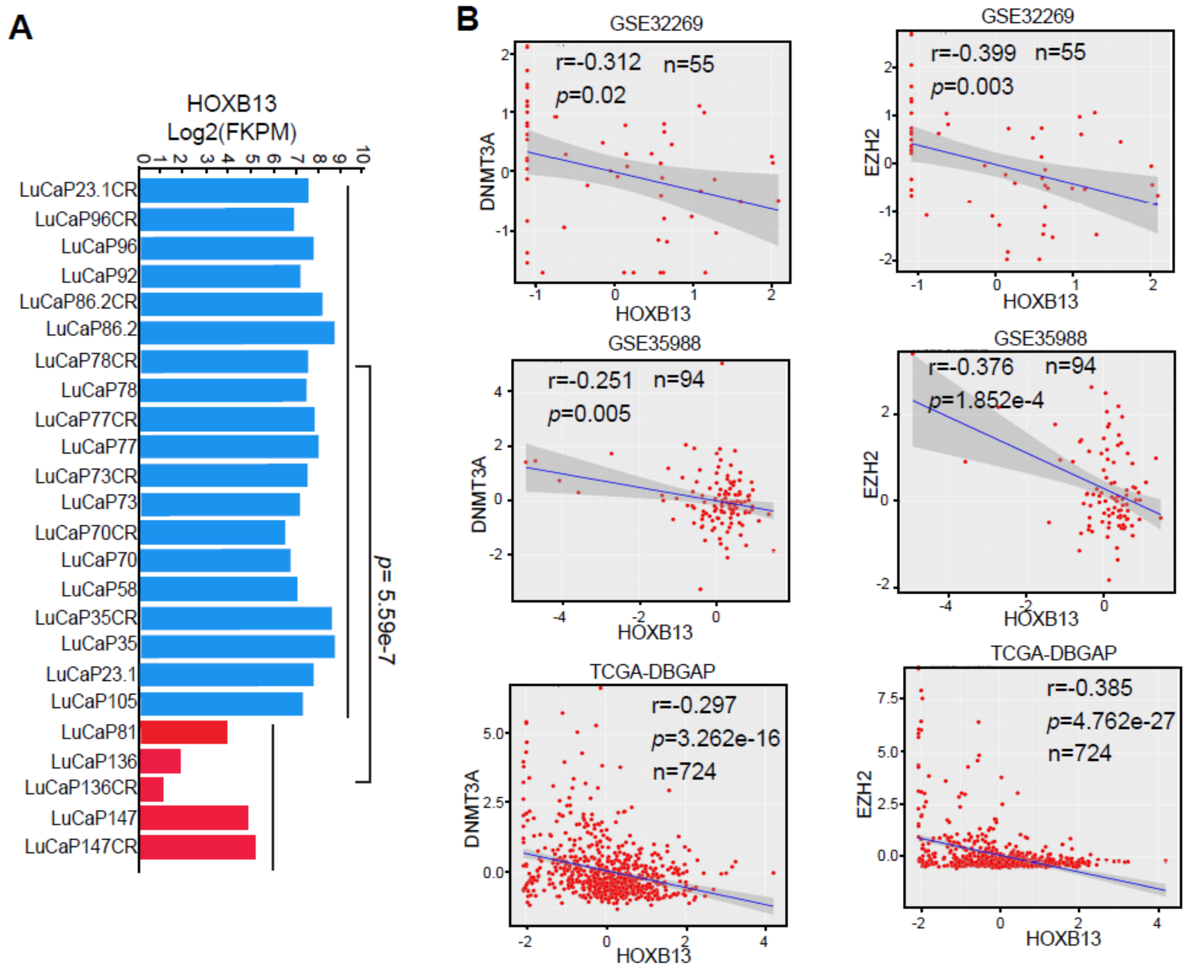


**Figure S5. HOXB13 is hypermethylated and down-regulated in CRPC.**

**A**. RNA expression of HOXB13 in mCRPC LuCaP PDXs by RNA-seq, Blue ones here are samples with higher RNA expression of HOXB13 and Red ones are samples with lower RNA expression of HOXB13 as show in Fig.5G.

**B**. Negative correlation between the mRNA of HOXB13 and DNMT3A (left) or EZH2 (right) in multiple publicly available PCa datasets.

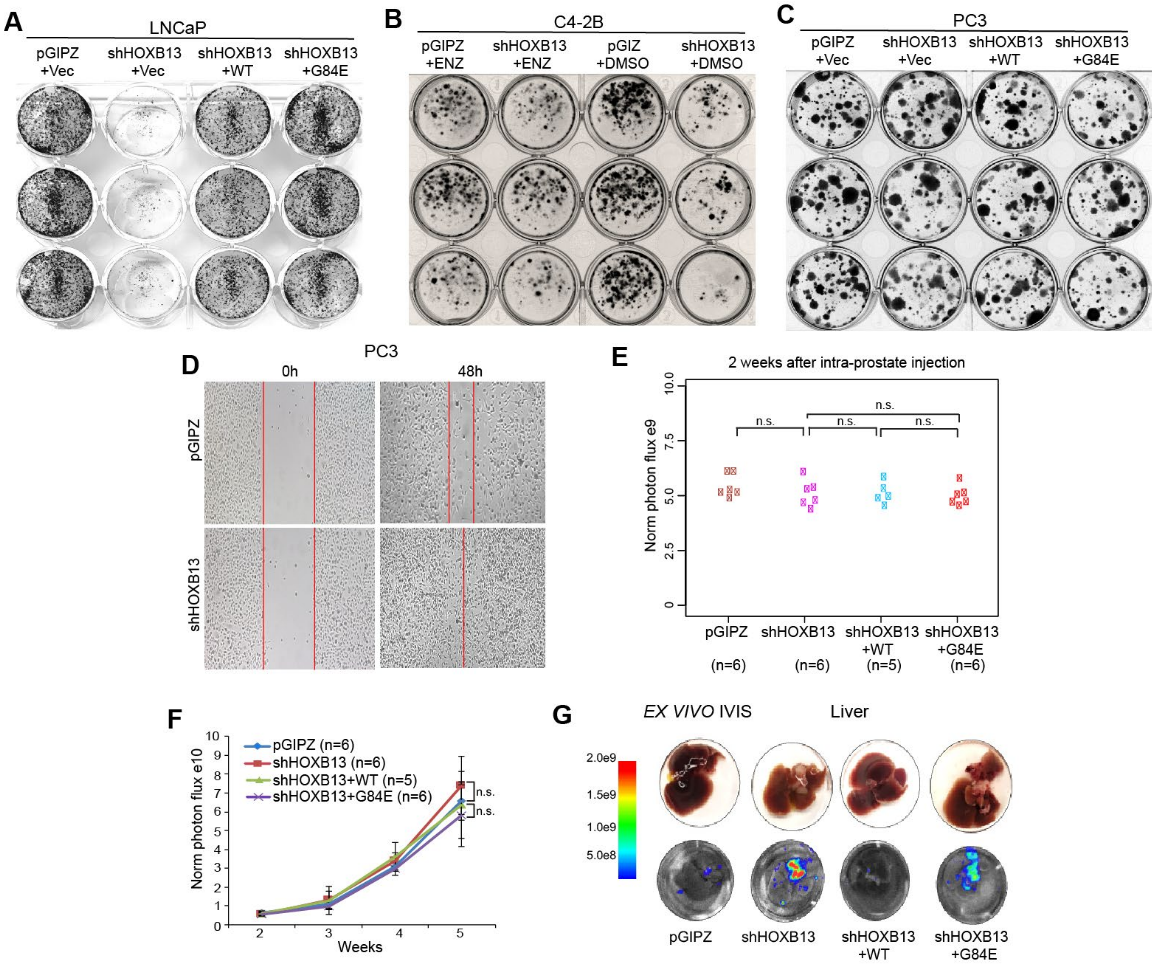


**Figure S6. HOXB13 loss promotes PCa cell motility *in vitro* and tumor metastasis *in vivo.***
**A-C.** Colony formation assays were performed in LNCaP, C4-2B, and PC-3 cells with indicated de-regulation of HOXB13 or treatment.
**D**. Cell migration assays of PC-3 cells with control or shHOXB13. Image shown were taken at 0 hour and 48 hours after a scratch was created on the cell monolayer.
**E**. Tumor volume after 2 weeks of intraprostatic inoculation of PC-3M cells. Luciferase-labeled PC-3M cells with control (pGIPZ), HOXB13 KD, HOXB13-KD with WT or G84E re-expression were inoculated orthotopically into the anterior prostates of nude SCID mice. A total of 23 mice were used with 5-6 mice/group. Tumor establishment rate was 100%. Tumor volume was measured by IVIS after two weeks of intra-prostate inoculation of PC-3M cells. Y-axis shows the normalized luciferase intensity. Statistical significance were evaluated by 1-way ANOVA and comparisons between 2 groups by Student's *t*-tests. *n.s.* not significant.
**F**. HOXB13 de-regulation did not affect PC-3M xenograft tumor growth. Tumor volume was measured every week by IVIS live mice imaging. Y-axis shows the normalized luciferase intensity. *n.s.* not significant as tested in **E**.
**G**. *Ex vivo* IVIS imaging of tumor cells metastasized to the liver. At the endpoint, mice were euthanized and livers were collected and IVIS was performed *ex vivo*. Data shown are representative *ex vivo* IVIS images of liver obtained from one mouse of each experiment group. Heatmap on the left indicates the intensity of IVIS signal.

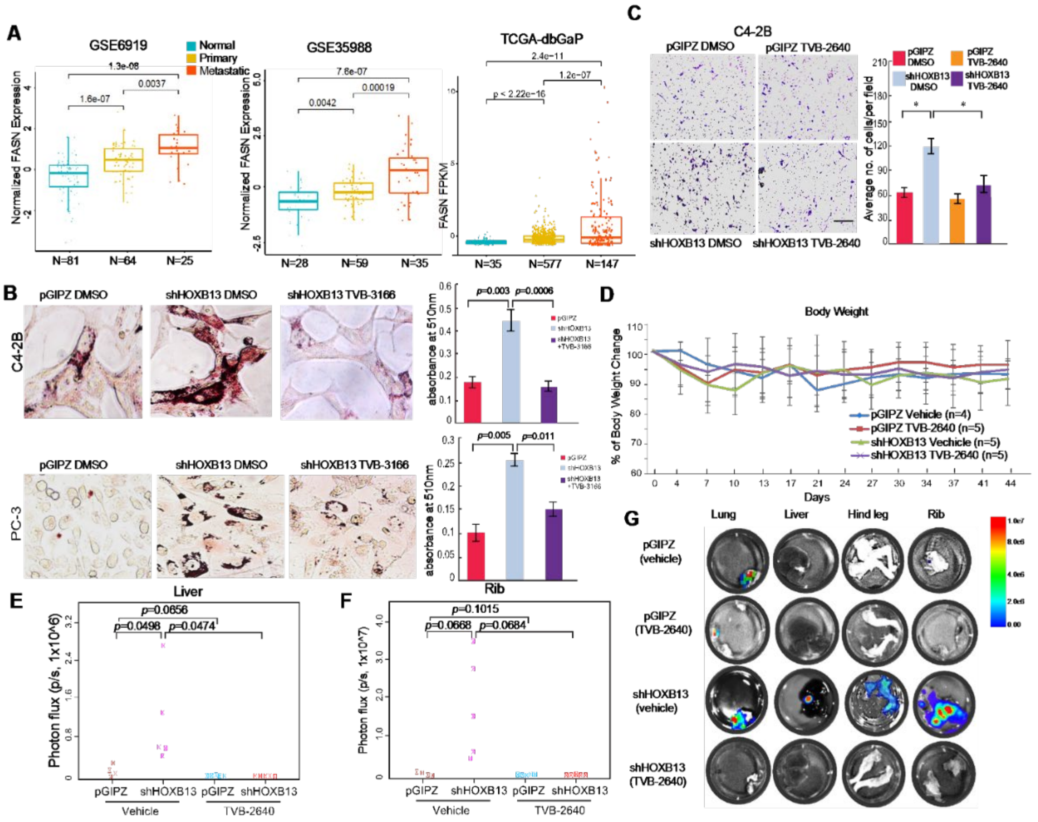


**Figure S7. Therapeutic targeting of HOXB13-low tumors with FASN inhibitors.**

**A.** FASN mRNA levels in publically available PCa gene expression profiling datasets.

**B**. Representative images of Oil Red O staining (left) and quantification (right) of lipid accumulation in HOXB13 KD C4-2B (upper) and PC-3 (lower) cells treated with DMSO or TVB-3166 for 3 days. *Scale bar* of 50 µm. Data shown are mean (±SEM) of triplicate wells. The *p* values shown are by two-tailed unpaired Student's *t*-test.

**C**. Cell invasion assays of control or HOXB13-KD C4-2B cells treated with DMSO or 2µM FASN inhibitor TVB-2640 for 3 days. Representative images are shown (left panels) and the number of invaded cells quantified (right panels). $^{**}p<0.01$, $^{*}p<0.05$, n.s. not significant by two-tailed, unpaired Student's t-test.

**D**. Body weight change analysis in vehicle and TVB-2640 treatment mice. Y axis shows percentage of body weight change.

**E-G**. *Ex Vivo* IVIS quantification of metastasis in liver and rib (**E-F**) and representative *ex vivo* IVIS images (**G**) of PC-3M tumor metastasis in lung, hind leg, liver and rib. At the endpoint, lung, hind leg, liver and rib were collected and performed *Ex vivo IVIS*. Y-axis shows the normalized luciferase intensity. Indicated *p* value were shown by two-tailed unpaired Student's t-test.

**Supplementary Table 1. Antibodies that were utilized in this study.**

| Antibody | Source | Catalog Number |
|---|---|---|
| Rabbit polyclonal anti-AR | Millipore | Cat# 06-680 |
| Rabbit polyclonal anti-AR | Santa Cruz Biotechnology | Cat# sc-816 |
| Rabbit polyclonal anti-HOXB13 | Santa Cruz Biotechnology | Cat# sc-66923 |
| Rabbit polyclonal anti-HOXB13 | Abcam | Cat# ab201682 |
| Rabbit monoclonal anti-HOXB13 | Cell Signaling Technology | Cat#90944S |
| Rabbit monoclonal anti-PSA | Cell Signaling Technology | Cat# 2475S |
| Rabbit polyclonal anti-H3K27Ac | Abcam | Cat# ab4729 |
| Rabbit polyclonal anti-FASN | Bethyl Laboratories | Cat#A301-323A-M |
| Mouse monoclonal anti-FASN | Santa Cruz Biotechnology | Cat#sc-48357 |
| Rabbit polyclonal anti-HDAC1 | Abcam | Cat#ab7028 |
| Rabbit polyclonal anti-HDAC3 | Abcam | Cat# ab7030 |
| Rabbit polyclonal anti-HA | Abcam | Cat# ab9110 |
| Mouse monoclonal anti-HA | Santa Cruz Biotechnology | Cat# sc-7392 |
| Rabbit polyclonal anti-Flag | Sigma | Cat#F7425 |
| Mouse monoclonal anti-Flag | Sigma | Cat# F1804 |
| Mouse monoclonal anti-Tubulin | Santa Cruz Biotechnology | Cat#sc-32293 |
| Rabbit Polyclonal anti-SREBF1 | Proteintech | Cat#14088-1-AP |
| Rabbit Polyclonal anti-SREBF2 | Proteintech | Cat#28212-1-AP |

**Supplementary Table 2. Oligonucleotides that were used in this study.**

| Name | Sequence (5′ to 3′) | Application |
|---|---|---|
| HOXB13-BamHI F | CGCGGATCCATGGAGCCCGGCAATTATGC | Cloning |
| HOXB13-XbaI R | CGCTCTAGAAGGGGTAGCGCTGTTCTTC | Cloning |
| HOXB13-70-284aa | CGCGGATCCCAGGGGACGTCCCCAGCT | Cloning |
| HOXB13-150-284aa | CGCGGATCCACTCTGGGTGCTCCTGGA | Cloning |
| HOXB13-210-284aa | CGCGGATCCTGCGCCTTTCGTCGCGG | Cloning |
| HOXB13-WFQ-3A F | TGCGAACCGCCGGGTCAAAGAG | Cloning |
| HOXB13-WFQ-3A R | GCCGCGATGGTAATCTGGCGCTC | Cloning |
| HOXB13-ΔMEIS F | ACTCTGGGTGCTCCTGGA | Cloning |
| HOXB13-ΔMEIS R | AGGGCATGGGTGGCATTG | Cloning |
| HOXB13-ΔHOX F | GAGAAGAAGGTTCTCGCCAAGG | Cloning |
| HOXB13-ΔHOX R | GCAGGCGTCAGGAGGGTG | Cloning |
| HDAC3-BamHI F | CGCGGATCCATGGCCAAGACCGTGGCCTA | Cloning |
| HDAC3-1-316aa XbaI | CGCTCTAGATTCTACCAGCAGCGATGTCTC | Cloning |
| HDAC3-1-180aa XbaI | CGCTCTAGAAATGTCGTTGACATAGCAGA | Cloning |
| HDAC3-1-120aa XbaI | CGCTCTAGAGTTGTTCAGCTGGGTTGCTC | Cloning |
| HDAC3-120-428aa | CGCGGATCCATGAAGATCTGTGATATTGCCAT | Cloning |
| HDAC3 XbaI R | CGCTCTAGAAATCTCCACATCGCTTTCCTTG | Cloning |
| NCOR1-ΔN1/2 F | GGTACTGCCAACACCTCA | Cloning |
| NCOR1-ΔN1/2 R | CTGCTGTGGTCGATAGTG | Cloning |

| | | |
|---|---|---|
| NCOR1-ΔN1/2/3 R | GGTGATGATCACGTCTATGAAG | Cloning |
| NCOR1-ΔDAD F | GCCCTCGTCAGAAGGAAT | Cloning |
| NCOR1-ΔDAD R | CATGTTAATGAACTTGACTCGTC | Cloning |
| PSA-qF | ACGCTGGACAGGGGGCAAAAG | RT-qPCR |
| PSA-qR | GGGCAGGGCACATGGTTCACT | RT-qPCR |
| FASN-qF | GAGTTCTGGGACAACCTCATC | RT-qPCR |
| FASN-qR | GAAGGAGGCATCAAACCTAGAC | RT-qPCR |
| SREBF1-qF | TGGGAGAGAGACGTGTACATAG | RT-qPCR |
| SREBF1-qR | CACTAGTCAGCACATCCATCAG | RT-qPCR |
| SREBF2-qF | GAGCACCAAGCACGGAGA | RT-qPCR |
| SREBF2-qR | GGGAGGAGAGGAAGGAGAGG | RT-qPCR |
| E2F1-qF | ACCCTGACCTGCTGCTCTT | RT-qPCR |
| E2F1-qR | ATGGTCAGTTTCCAGGTCCA | RT-qPCR |
| MYBL2-qF | CTCAACCCTGAGGTGAAGAAG | RT-qPCR |
| MYBL2-qR | TCACAGCATTGTCTGTCCTC | RT-qPCR |
| FOXM1-qF | TCTGGAGGGTCCACACTT | RT-qPCR |
| FOXM1-qR | CCTCCTCTGATGTTTCACTTGG | RT-qPCR |
| CDCA2-qF | GACAGAGCATGTGCAGTTGAA | RT-qPCR |
| CDCA2-qR | TGAGCTCTGAAAGGGGAAGA | RT-qPCR |
| PSA-AF | GCCTGGATCTGAGAGAGATATCATC | ChIP-qPCR |
| PSA-AR | ACACCTTTTTTTTTCTGGATTGTTG | ChIP-qPCR |
| FASN-AF | CAGAAGAGTAAACGCAGGAGAA | ChIP-qPCR |
| FASN-AR | CCTCACTTTAGGACCAGGAAAC | ChIP-qPCR |
| SREBF1-AF | GCTGATGGATGTGCTGACTA | ChIP-qPCR |
| SREBF1-AR | CACTCCTCCCACTAACAAACA | ChIP-qPCR |
| SREBF2-AF | TATCGAAGGGCTTGCAAACTA | ChIP-qPCR |
| SREBF2-AR | CTAGATCCTGAACCCAACAGAC | ChIP-qPCR |
